# Kinematic fields of the visuo-hippocampal circuit

**DOI:** 10.64898/2025.12.30.697009

**Authors:** Chinmay S. Purandare, Mayank R. Mehta

## Abstract

The structure of focused receptive fields in retinotopic and allocentric space in the visual cortex and hippocampus is studied extensively to understand cognitive functions such as stimulus tracking and path-integration, respectively. These functions require accurate tracking of running speed and acceleration. While speed dependence in these areas is well-studied, the acceleration dependence has received little attention, and the joint effect of speed-acceleration is unknown. Hence, we assessed the joint influence of these kinematic variables by computing receptive fields in a novel, 2D phase space of speed-acceleration using the Allen Brain Visual Observatory data. Head-fixed mice ran spontaneously on a running wheel next to a monocular gray screen ensuring virtually no changes in the retinotopic or allocentric space. Remarkably, about half (44.3%, 6590/14862) of neurons in the visuo-hippocampal circuit (from LGN to subiculum) were significantly modulated by a specific combination of speed and acceleration to form receptive fields in the speed-acceleration phase space (called kinematic space). The prevalence of kinematic tuning is comparable to the fraction of place cells in this part of the mouse hippocampus, and the fraction of visually tuned neurons in higher visual areas. The kinematic field size (∼30% of the sampled space) was also comparable to the place field size. Kinematic fields spanned the entire space but preferred either low-speed, high-acceleration or the high-speed, low-acceleration segments. Although pupil size too varied spontaneously, the kinematic tuning exceeded pupil size tuning in all brain areas. Surprisingly, pupil size modulated the activity of ∼20% of hippocampal neurons, which was comparable to that in LGN and visual cortical areas. Thus, all visuo-hippocampal areas have focused kinematic fields that are three times more prevalent than the pupil size fields in the absence of any vestibular, visual or spatial stimuli. This kinematic phase space could interface with retinotopic and allocentric spaces to guide natural behaviors involving movement.

## Introduction

Self-motion modulates neural activity in many brain areas^1–17^. In addition to visual stimulus motion^18–25^, visual cortical neurons are modulated by running speed too^3,7,26–28^. The interaction between self-and stimulus-motion shapes visual cortical selectivity and mismatch encoding^26,29,30^. At the other end of the visual hierarchy^31^ is the hippocampus which is thought to contain an abstract cognitive map of allocentric space^32^, generated via path, i.e. self-motion integration^33–36^. Consistently, running speed modulates the hippocampal^10,17,37,38^, entorhinal^14,17^, septal^37,39^, and subicular activity^11^. Further, running speed modulates the frequency^40^, amplitude and theta-phase of hippocampal gamma oscillations^41^, and the firing rate of hippocampal inhibitory interneurons^13,41,42^. Rodent hippocampal theta oscillations are also modulated by the running speed^43–47^ and acceleration^48^ (but see ref.^43^) potentially through the lateral septum input^39^.

### Kinematic phase space hypothesis

Visual cortical responses have been studied extensively by parametrizing neural activity in the X-Y (azimuth vs. elevation), retinotopic cartesian space^49–51^. Similarly, hippocampal responses have been studied as selectivity to the animals’ position in the X-Y cartesian space^32,52–54^, as well as other one^47^ or three-dimensional^55–58^ spaces. However, the movement of any system or an organism in space can also be quantified in a phase space - consisting of a variable *q* (e.g., position), and its derivative (*dq/dt*, e.g., velocity). In the Lagrangian or Hamiltonian formulation of mechanics, position and velocity are treated as independent variables that form a phase space where the dynamics is fully described^59,60^, even though these two variables are often mathematically linked, e.g. in the case of a pendulum or a simple harmonic oscillator, where each position has a unique speed.

However, organisms need not be born with the knowledge of the structure of space and need to learn this from experience. In this regard, the running speed (*s*) is a Euclidean variable that all animals experience in their egocentric coordinate frame. Unlike velocity, speed makes no assumptions about the dimension or structure of the embedding space. Further, running speed (*s*) and its temporal derivative, running acceleration (*ds/dt*), form a two-dimensional, kinematic phase space that can describe kinematics of ongoing behavior, without assuming anything about embedding space. Thus, the same self-motion metric is applicable in one, two, three, or even other-dimensional natural foraging, providing a general yet complete description of self-motion, in a behaviorally relevant fashion.

Such kinematic phase space is indeed plausible because, more generally, the geometry of dynamics is accurately described using other types of phase spaces too^61^. Similar to a simple harmonic oscillator and other physical systems, organisms may experience many, but not all possible combinations of speed and acceleration. On the other hand, unlike simple particles, speed and acceleration of organism would be accompanied by a complex array of muscular movements. Despite this, we commonly perceive just speed and acceleration. Hence, we hypothesize that visuo-hippocampal neurons, in charge of perceiving space, will encode this kinematic phase space by responding selectively to specific regions of the speed-acceleration space.

During natural behaviors, self-motion causes changes in visual and other stimuli, making it difficult to isolate the contribution of self-motion, let alone self-acceleration, or the joint effect of running speed and acceleration. Here we show that there is a circuit-wide neural code for the speed-acceleration phase space across all the seven visuo-hippocampal brain areas investigated, from LGN to subiculum. Pupil size too had a significantly greater than chance effect on neural activity in all parts of this circuit. The kinematic modulation was more than twice as prevalent in the entire circuit as pupillary tuning. The kinematic modulation resulted in focused fields in the speed-acceleration space. Kinematic fields preferred either the low-speed high acceleration quadrant or high-speed low acceleration segment of the kinematic space. Qualitatively similar results were found in putative interneurons across all brain regions. We hypothesize that these are universal, cortex-wide interoceptive or proprioceptive signals for self-motion.

## Results

Head-fixed mice were placed on an oblique running-wheel (Fig. 1A, see *Methods*). A monocular gray screen spanning the entire left visual field was presented while neural signals were measured from the contralateral hemisphere. No visual cues were shown to the ipsilateral eye. Mice ran spontaneously, without any reward or task demand and their running did not result in changes in the visual input, allowing us to estimate the contribution of pure self-motion cues on neural activity. To focus on the contribution of self-motion, without the effect of brain-state changes, data from long bouts of immobility were discarded and only the data from bouts of locomotion (speeds > 2cm/sec) were used, including up to 5 seconds before and after the running bout (see *Methods*). We first focused on the broad spiking, putatively excitatory neurons from the thalamo-cortical (Lateral geniculate nucleus, LGN (n=268), primary visual cortex V1 (1726), higher visual areas (antero-medial and posterior-medial) AM&PM (2226), and the (intermediate and ventral) hippocampus (dentate gyrus DG (1037), CA3 (1377), CA1 (7456), & Subiculum SUB (708)) from 18 mice, one session per animal^62^.

**Figure 1.**
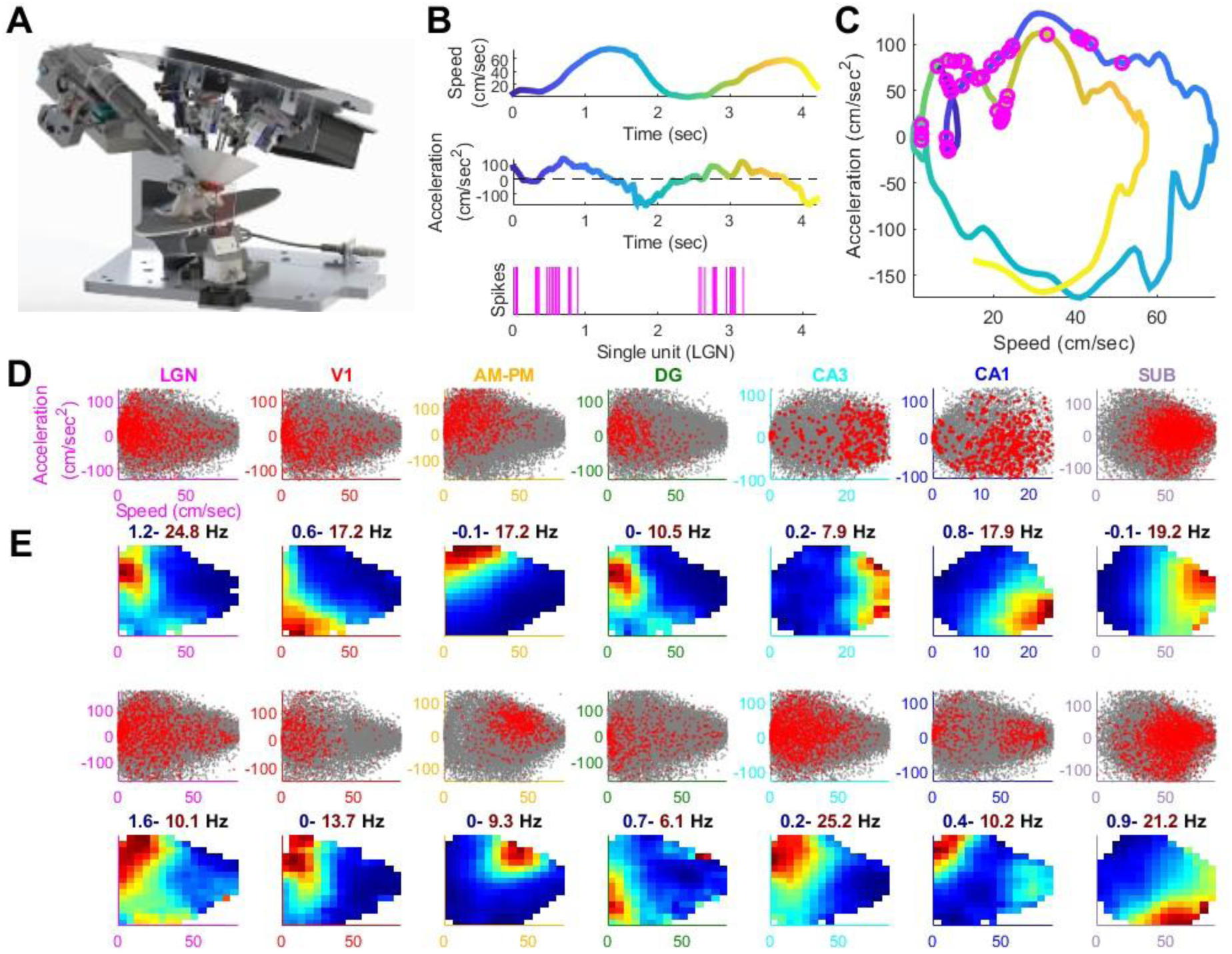
Kinematic fields in the visuo-hippocampal circuit: **(A)** Experimental setup with an oblique running wheel on which head-fixed mice ran spontaneously. There was a large gray screen (not shown) to the left of the mice and the room was dark. Neural data were recorded from the contralateral hemisphere. **(B)** (Top) Running speed color coded as a function of time. (Bottom) Spikes from a representative LGN cell that fired (magenta lines) during increasing speed (acceleration), but not decreasing speed (deceleration). **(C)** Data from (B) transformed into the kinematic space of speed-acceleration with overlaid spikes (magenta circles). **(D)** Two representative cells each from seven brain regions showing elevated firing in a restricted region of the 2D kinematic space of speed (x-axis) and acceleration (y-axis). Gray dots denote instantaneous speed-acceleration, red dots denote spikes. **(E)** The resulting firing rate maps show clear kinematic fields. The firing rate modulation range is indicated above the kinematic rate maps. Bottom two rows are another set of example cells, in same format as (D) and (E).

### Kinematic fields in all seven brain regions

To determine the joint effect of running speed and acceleration on neural firing, we computed tuning curves in a 2D kinematic space, comprised of instantaneous speed and acceleration (Fig. 1D-E). Many neurons in all brain regions showed elevated firing in localized regions, termed kinematic fields, of this two-dimensional kinematic space of speed-acceleration (Fig. 1D-E, Fig. S1). Instead of purely speed or acceleration tuning (which would manifest as vertical or horizontal bands of elevated firing rates respectively), neural responses were jointly modulated by both speed and acceleration, resulting in kinematic fields, showing joint encoding of a specific combination of speed-acceleration. This joint modulation in the speed-acceleration 2D space was called kinematic tuning. The statistical significance of kinematic tuning was quantified by bootstrapping methods^63,64^ (see *Methods*). At least 30% of the broad spiking neurons showed significant (z>2, equivalent to a *p*<0.023 in a one-sided Student’s t-test) kinematic tuning in each of the seven brain areas, with CA3 showing the smallest prevalence (31%) and the higher visual areas, AM & PM showing the largest prevalence (61%) (Fig. 2A).

**Figure 2.**
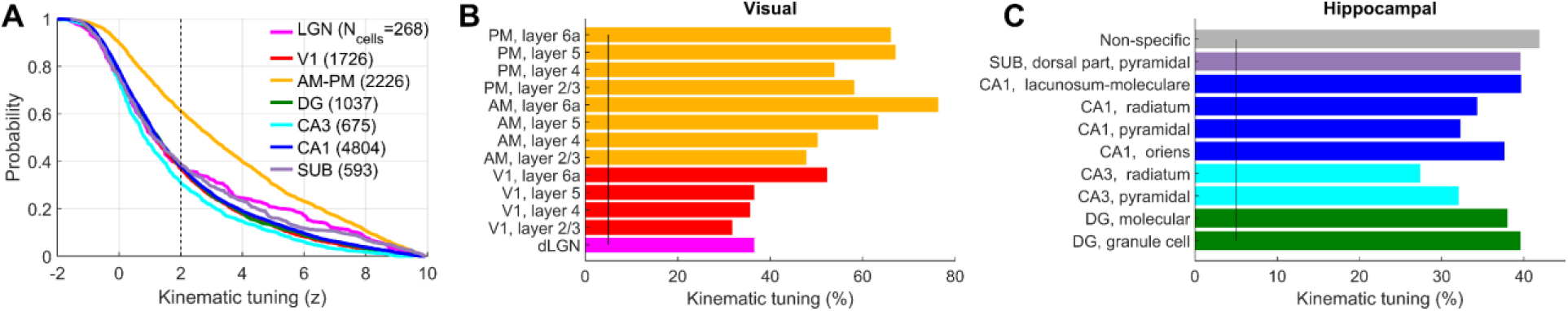
At least one third of neurons in all brain areas and subregions show significant kinematic tuning. **(A)** Cumulative histogram of kinematic tuning strength for all broad spiking neurons (see *Methods*). All regions had significantly higher prevalence than expected by chance (∼2.5%), based on a z-scored sparsity >2 threshold (dashed vertical line). The number of neurons used is indicated with the legend. The highest prevalence of kinematic tuning was found in higher visual areas, AM-PM (61.1%) and the lowest prevalence was seen in CA3 (31.0%). **(B)** Sublayer dependence of kinematic tuning. The unit location was identified using anatomical assignment to the average brain template (see *Methods*). Only those sublayers with at least 100 broad spiking neurons were used for statistical rigor. All subregions had significantly greater (Kolmogorov-Smirnov, KS-test, here and subsequently, unless mentioned otherwise, *p*<3.9×10^-11^) kinematic tuning than expected by chance (∼2.5%, black vertical line) with the highest prevalence seen in layer 6a in V1 (52.3%) and in higher visual areas (AM-76.3%, PM-66.0%, but PM-layer 5 was slightly higher at 67.1%). Layer 2/3 had the lowest prevalence in V1 (31.7%) and similarly for AM-PM. Across the visual cortical regions, deeper layers had significantly larger kinematic tuning compared to the superficial layers (Layers 5&6 vs Layers2/3 & 4, *p*<5.4×10^-3^). **(C)** Similar to (B), all hippocampal subregions had significant kinematic tuning, with the largest prevalence in CA1 lacunosum-moleculare (39.6%) and least in CA3 radiatum (27.4%). Unlike visual areas, kinematic tuning rates were relatively independent of hippocampal subregions and layers.

### Cortical layer-dependent but hippocampal region-independent kinematic tuning

To understand the finer anatomical organization of kinematic tuning, we leveraged the high-density recording capabilities of *Neuropixel* probes combined with registration of recording sites to a common anatomical coordinate framework^65^, to compute the prevalence of kinematic tuning in different subregions of visual thalamus (LGN), cortex (Fig. 2B) and the hippocampus (Fig. 2C). All sublayers of primary (V1) and higher visual areas (AM-PM) showed significant tuning.

The kinematic tuning in L4 of V1 was comparable to LGN (36% each). However, in V1, kinematic tuning was 1.6*x* more prevalent in the deeper layers especially layer 6 (52.3%), compared to superficial layers 2/3 and 4 (31.7%). A similar, but weaker layer-dependence was found in AM-PM. Across the three visual cortical regions (V1, AM and PM), deeper layers had greater (1.21*x*) kinematic tuning compared to the superficial layers (Layers 5&6 vs Layers 2/3&4).

Kinematic tuning prevalence showed much smaller variation across different parts and subregions of the hippocampus (range of 27.4 to 41.8%) compared to visual areas (31.7 to 76.3%). The highest prevalence of kinematic tuning was in the deep layer of CA1 (lacunosum-moleculare) and least in CA3 radiatum. Thus, kinematic tuning was significantly above chance levels in 27.4% to 76.3% (mean value of 45.1%) of neurons in all sub-layers of the thalamo-cortical-hippocampal visual circuit. The differences in firing rates^52,66–69^ and sampling across sessions could affect the differences in percentage tuned cells, however the presence of tuning across layers and brain regions demonstrates the robustness of these results.

### Kinematic fields accumulate in regions of low-speed, high-acceleration or high speed

Although the kinematic field-peaks spanned the entire speed-acceleration space, they largely over-represented the edges of this space (Fig. 3A-B, see *Methods*). Kinematic field peaks accumulated in the region of low-speed and high-acceleration and the region of high-speed and low-acceleration (Fig. 3C). Very few kinematic field-peaks were found in the middle of the kinematic space, e.g. regions of intermediate speeds and acceleration. Further, the regions of high deceleration were also under-represented. The kinematic field-density distributions were remarkably similar across brain regions (Fig. 3F), with the least similarity between LGN and AM-PM.

**Figure 3.**
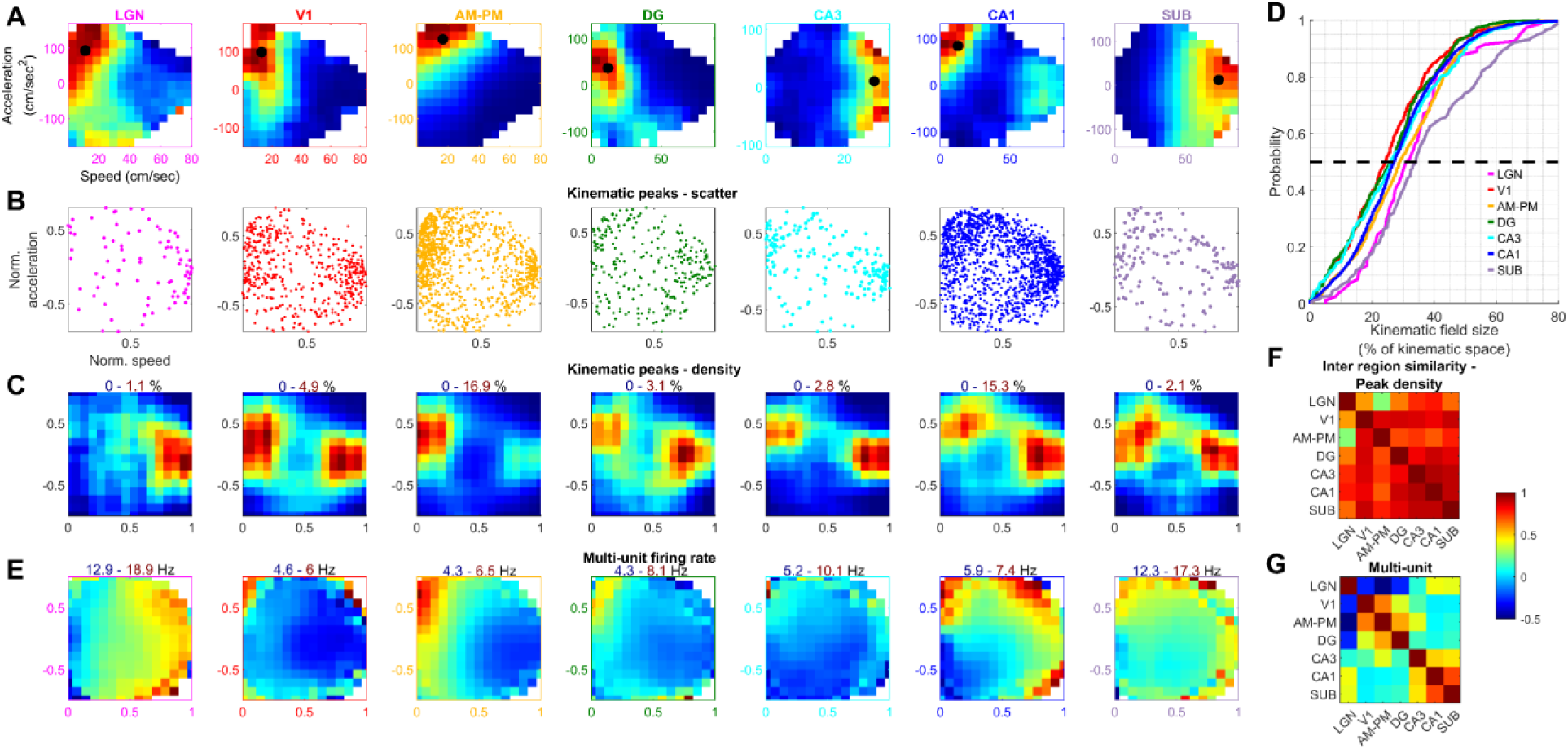
Kinematic fields overrepresented specific combinations of speed and acceleration in each brain area. **(A)** Representative broad-spiking cells’ kinematic responses from each brain region, same as Fig. 1, overlaid with a black dot indicating peak response location (see Methods). (**B)** Each dot depicts the kinematic field peak location of an individual, significantly tuned, broad spiking cell. **(C)** Density plot corresponding to (B). Density plot (color indicates percentage of data) of the distribution of kinematic field locations across all tuned cells from that brain region. In all brain regions, low speed-acceleration was preferred (31.3% to 46.2% of peaks in the top-left quadrant), except LGN (24.4%), which preferred high speed (61% of peaks to the right). Low-speed deceleration (bottom-left) was least preferred by all brain regions (12.8% in CA3 to 21.1% in DG, compared to 25% chance for each quadrant) except V1 and AM-PM. The density range is indicated above each color plot. **(D)** Cumulative histogram of the kinematic field sizes, computed as the percentage of the explored kinematic space. Black dotted line indicates median crossing. The kinematic field size distributions (colored lines) were similar across brain regions with the mean field sizes ranging from (mean±s.e.m., here and subsequently, 25.2±0.7%) in V1 to (37.1±1.4%) in subiculum. **(E)** Average spiking activity across all broad spiking kinematic tuned cells from a given brain region across all mice and sessions. Since the range of speed-acceleration varied across sessions, the range was normalized between 0-1 for speed and-1 to +1 for acceleration corresponding to the lowest and largest values in that session. The range of firing rates for each color plot is indicated above the plot. The mean firing rates were the highest for LGN (16.0±0.06Hz) and subiculum (14.8±0.04Hz), nearly twice as large as the mean firing rates in all other brain regions (6.2±0.2Hz). **(F)** Similarity between kinematic field density maps across different brain regions, defined as the correlation coefficient of the density plots in (C). The kinematic field density maps between LGN and AM-PM were the least similar (Pearson’s correlation coefficient, here and subsequently, r=+0.26, p=3.2×10^-5^) but they were most similar across CA3, CA1 and subiculum (r>+0.9, p<2×10^-96^). **(G)** Similar to (F), for the multi-unit kinematic rate maps across brain regions in (E). Same colormap as (F). LGN multi-unit responses were significantly negatively correlated with those for V1, AM-PM and DG (r<-0.27, p<8×10^-5^). LGN-CA1, LGN-SUB, AM&PM-DG, CA3-CA1, CA3-SUB and CA1-SUB brain region pairs were significantly positively correlated (r>+0.39, p<4.3×10^-9^), and other brain region pairs were not significantly correlated (p>0.05).

Next, we used standard techniques to define the borders and size of kinematic fields (see *Methods*), i.e. to identify the restricted region of the kinematic space where a tuned neuron fired at an elevated rate. On average, each kinematic field covered a third (29.3%) of kinematic space, based on a threshold from the shuffle data (Fig. 3D, see *Methods*). This degree of specificity, i.e. the kinematic field size as a fraction of explored space is similar to visual motion receptive fields in CA1^63^, distance tuned cells in CA1^47,70,71^ and place fields in many parts of the hippocampal formation during spatial exploration^52^. On the other hand, the kinematic fields were larger than the visual cortical retinotopic receptive fields in the nasal or foveal areas^62^. However, the precise size of the kinematic fields, as well as the receptive fields in other experiments, depend on the definition of the field boundary, and alternate choices can make the fields bigger or smaller.

The above method relies on the definition of kinematic field boundary. Another way to quantify ensemble wide kinematic responses, independent of the kinematic field identification, is to compute the average response of the ensemble of all tuned cells (multi-unit firing activity, see *Methods*). In cortex and hippocampus, the neural ensembles preferred low speed-acceleration epochs, corresponding to locomotion initiation (Fig. 3E), consistent with the above findings and with recent reports^27^. Further, similar to the kinematic field-density results, the lowest ensemble-wide activity occurred at intermediate speeds in all brain regions. Thus, the multi-unit responses matched the kinematic field density in many ways, but there were considerable differences as well. For example, LGN, CA1 and subiculum preferred high speed over low speeds, whereas visual cortical regions as well as dentate gyrus and CA3 reduced activity at high speeds. AM-PM kinematic fields preferred high speed, whereas the multi-unit response showed a reduction in activity at high speeds. Similar differences existed in dentate gyrus and to a smaller extent in V1, which could be in part due to the differences in kinematic field properties across regions. The ensemble responses were most similar between CA1-subiculum and least between LGN and AM-PM (Fig. 3G), similar to the ensemble level comparison of kinematic peaks.

### Kinematic tuning is diminished but persists in the presence of a movie

Does self-motion tuning persist in the presence of moving visual cues? We computed the kinematic tuning while a black-and-white, silent movie clip^62,72^ covering the entire hemi-field was presented (See *Methods*). Similar to the gray screen condition, the locomotor behavior had neither a direct impact on the visual cues, nor on rewards. Many kinematically selective neurons were found in this case as well (Fig. S2). Across all the brain regions, kinematic tuning prevalence during movie presentation was above chance, but slightly smaller than that in gray screen presentation. Hence, the encoding of self-motion was not abolished when dynamic, spatio-temporal visual cues, which have been shown to generate selective activity in the entire visuo-hippocampal circuit^72^, were presented. Surprisingly, the kinematic fields of the same cell during the blank screen versus movie conditions were poorly correlated (average *r*=0.34) indicating a non-trivial interaction of self-and stimulus-motion on neural coding.

### Putative inhibitory interneurons have similar kinematic fields as broad spiking neurons

Speed tuning has been widely studied in the hippocampal interneurons^12,13,41^ (which have narrow spike waveforms). The interneuron firing rate is commonly believed to increase with running speed^13,14,41,42^. This has also been reported for pyramidal cells, which are putatively excitatory, on a running wheel^10^. However, a recent study of CA1 showed that only interneurons encoded running speed, whereas excitatory neurons were merely modulated by but did not encode running speed^73^. To address these differences, we conducted the above analysis (which was restricted only to excitatory, broad spiking neurons) for the narrow spiking interneurons (see *Methods*). Interneurons in all brain regions showed significant kinematic tuning (Fig. 4A-B). Unlike previous studies, most kinematic tuning properties were quite similar between the broad and narrow spiking cells, including layer dependence in visuo-hippocampal regions (Fig. 4C-D), size of kinematic fields (29.5±1.3% in V1 to 37.4±1.9% in CA3, Fig. 4E) and bias for acceleration (Fig. 4F-G).

**Figure 4.**
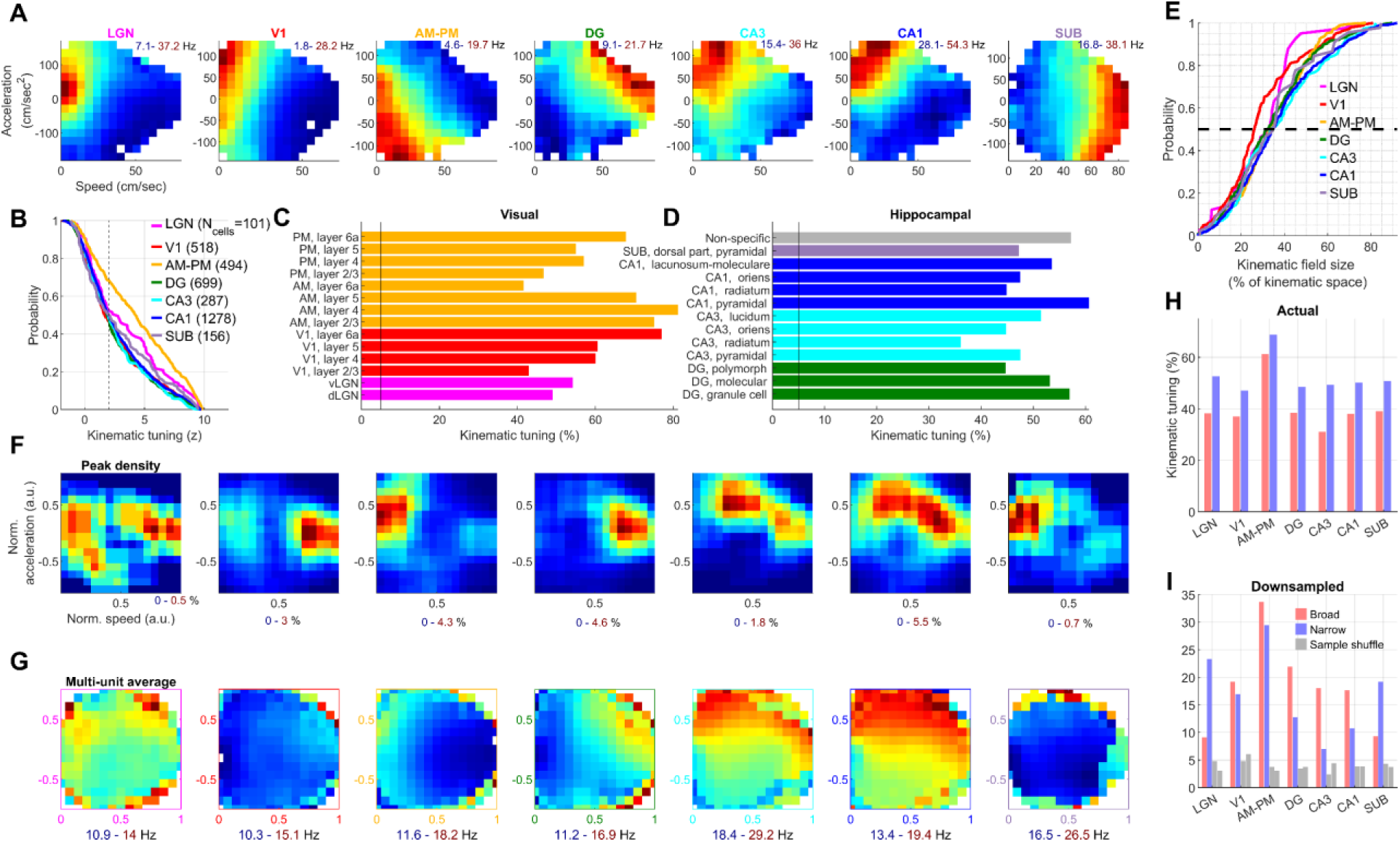
Similar kinematic tuning properties of inhibitory cells as of the excitatory cells. **(A)** Similar to Fig. 1E, but for narrow spiking cells showing robust kinematic fields. Note the higher firing rates (inset, top-right) than for broad spiking neurons in Fig. 1E. **(B)** Cumulative histogram showing similar prevalence of kinematic tuning as Fig. 2A with the highest tuning prevalence in AM-PM (higher visual areas) at 68.6% and lowest in DG at 48.4%. **(C)** Similar to the excitatory cells, all visual subregions had significantly higher kinematic tuning than expected by chance (∼2.5%, black dotted line), with the highest prevalence seen in layer 4 in AM (81.1%) and least in layer 6a of AM (41.5%). Similar to the broad spiking units, only sublayers with at least 30 narrow spike waveform cells were used to ensure sufficient statistical power. **(D)** Similar to (C), significant kinematic tuning was seen in all hippocampal subregions’ interneurons, with the largest prevalence in CA1 pyramidal cell body layer (60.7%) and least in CA3 radiatum (36.1%). **(E)** Similar to Fig. 3D, kinematic field sizes for narrow spiking cells varied over a large range, with the mean field sizes ranging from (29.5±1.3%) in V1 to (37.4±1.9%) in CA3. **(F)** Density plot of kinematic field peaks, similar to Fig. 3C, were biased towards acceleration compared to deceleration for all brain regions as well as epochs of highest speed. Top-right (low speed-acceleration) was the most preferred kinematic behavior for LGN, AM-PM, CA3, CA1 and SUB, whereas high speed was the most preferred for DG and V1. **(G)** Similar to Fig. 3E for narrow spiking cells showing ensemble wide biases. **(H)** Fraction of kinematically tuned cells was significantly (*p*<0.03) higher for narrow than broad spiking units in all brain regions. **(I)** The firing rates of neurons influence the significance levels of tuning. To address this, random spikes were deleted from each unit to have an effective firing rate of 1Hz (see *Methods*). Cells below 1Hz firing rate were not used for this analysis. Except for LGN and subiculum, all brain regions showed greater kinematic tuning prevalence in broad spiking cells based on this comparison, and the tuning levels were not significantly different for V1 and AM-PM. Gray bars show sample shuffle data. The overall reduced levels of kinematic tuning in all populations here are due to the removal of spikes, compared to Fig. 2A or 4B.

A few differences were found as well - CA3 and CA1 interneurons preferred intermediate-speed, high-acceleration region of the kinematic space, unlike the excitatory neurons. Similarly, V1 interneurons differed from their respective excitatory cells since they did not have as strong of a preference for low-speed high-acceleration. Similar results were found using alternate definitions of cell-classification e.g., using mean firing rate or complex spiking index-based metrics^63,74,75^ (see *Methods*, Fig. S3). The similarity of kinematic fields in broad and narrow spiking cells is in stark contrast to hippocampal spatial selectivity where place fields are sharp in excitatory cells but diffuse in interneurons^12,53,76–78^.

Prevalence of kinematic tuning was higher for narrow spiking cells in all brain regions (Fig. 4H). However, the conventional methods of neural tuning are strongly influenced by the mean firing rates^64,79^. Consistent with previous studies, inhibitory interneurons had higher firing rates than broad spiking neurons (5.4±0.06Hz for broad vs. 12.2±0.2Hz for narrow spiking neurons). Hence, we did additional analysis to remove the effect of neural firing rates across cells (see *Methods*).

This revealed that putative excitatory cells had a greater (1.1*x* to 2.6*x*) prevalence of kinematic tuning than inhibitory neurons across all brain regions, except LGN and subiculum (Fig 4I). Hence, contrary to previous studies, the kinematic variables of speed and acceleration affect inhibitory neurons across all brain regions in the visuo-hippocampal hierarchy, similar to the excitatory neurons but their apparent stronger encoding of kinematic space can be partially attributed to their higher firing rates. The differences between these results and previous studies could arise because of these improved analysis methods and because in our experiments the kinematic behavior did not cause any changes in the visual cues, which was not the case in other studies.

### Pupil dilation tuning in the visuo-hippocampal circuit

Similar to self-motion, pupil dilation also changes the visual input. Recent work has shown that along with self-motion^3,7,26^, visual responses are modulated by arousal^80^, which is also quantified by pupil dilation, and is believed to modulate visual cortical activity independent of running speed^80^. But the hippocampal tuning for pupil size is unknown. Surprisingly, many neurons in the visual (Fig. 5A) as well as hippocampal regions (Fig. 5B) showed ramp-like firing activity, increasing firing rate with increasing pupil dilation. Other neurons showed the opposite effect, with decreasing firing with increasing pupil size (ramp-like preference for pupil constriction).

**Figure 5.**
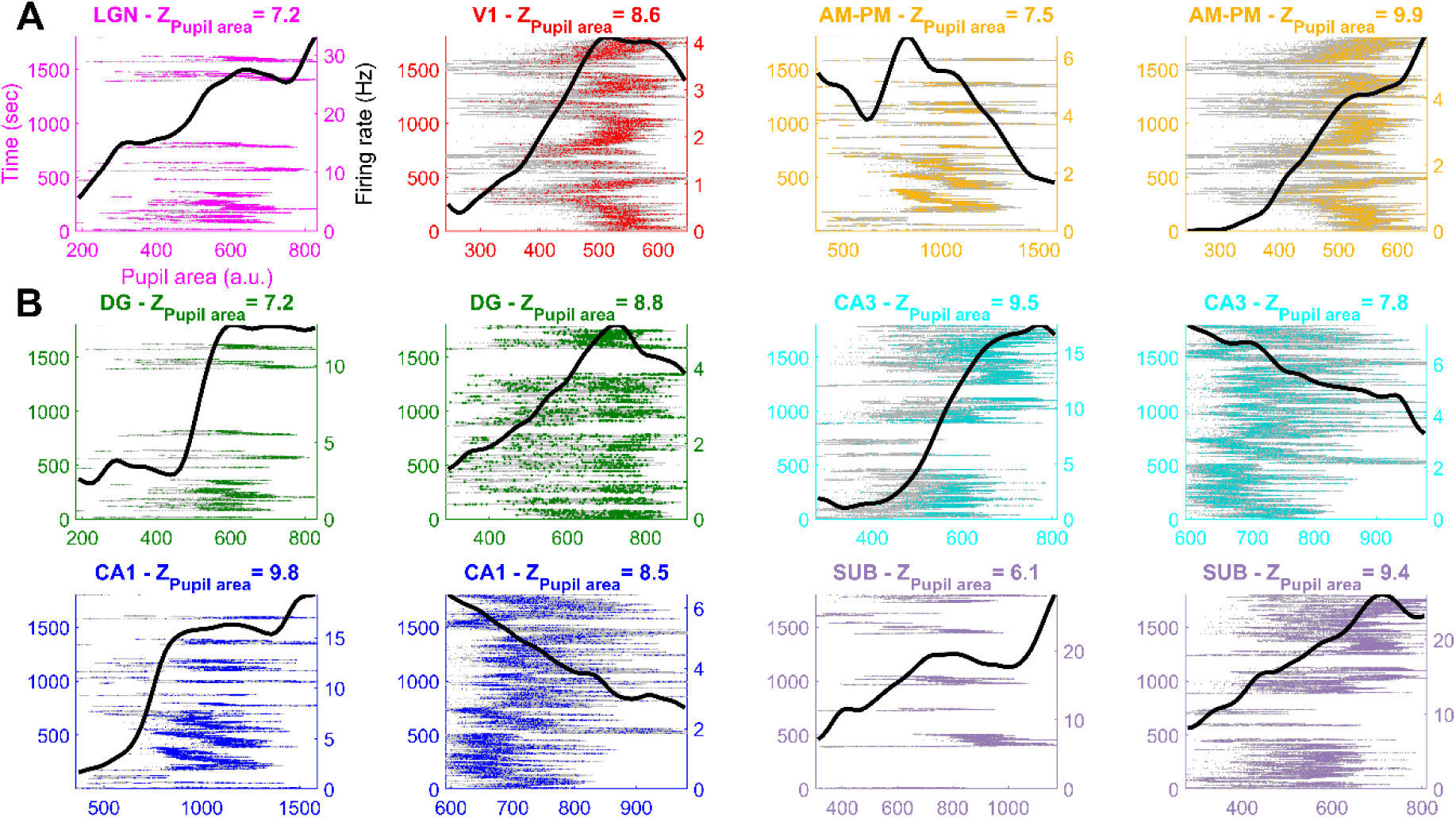
Pupil area tuned cells across the visuo-hippocampal circuit. **(A)** Four representative cells from visual regions show modulation in their mean firing rate responses (black traces) to pupil area. Gray lines indicate occupancy, colored dots indicate spikes from a neuron. Blank areas indicate epochs of long periods of immobility, and those data were excluded to rule out behavioral state changes (see *Methods*). Resultant strength of tuning, quantified as a z-score, is indicated above. **(B)** Similar to (A), for eight hippocampal neurons.

### Isolating kinematic and pupillary dynamics contributions using GLM

Does kinematic tuning persist independently of pupillary tuning? Firstly, we did several tests to rule out the possibility that kinematic tuning is an artefact of pupil area modulation of neural firing. Locomotion onset is often related with changes in arousal states, which are typically quantified by pupil dilation^81,82^, which in turn are hypothesized to alter neural firing in the visual areas^80^. We found that running speed changes were positively but weakly correlated with pupil size changes, during acceleration as well as deceleration (Fig. S4). Using the average neural response to pupil area, we predicted the kinematic tuning during running epochs (see *Methods*).

These were not significantly correlated with the observed kinematic tuning curves (Fig. S5). Hence, kinematic tuning is not entirely an artefact of pupillary dynamics.

To dissect the independent contributions of kinematic and pupillary dynamics on spiking, we employed a Generalized Linear Model (GLM, see *Methods*), which can separate simultaneous contributions of correlated variables^63,70,79^. The kinematic fields obtained using the GLM methods (Fig. 6A) were quite similar to those obtained using binning method (Fig. 1) in many ways, e.g. both methods revealed comparable fractions of tuned neurons and their nature of the tuning curves in all brain regions (Fig. 6B). This is consistent with the observation that while the pupillary and kinematic variables are correlated, the correlation is quite weak (∼0.15).

**Figure 6.**
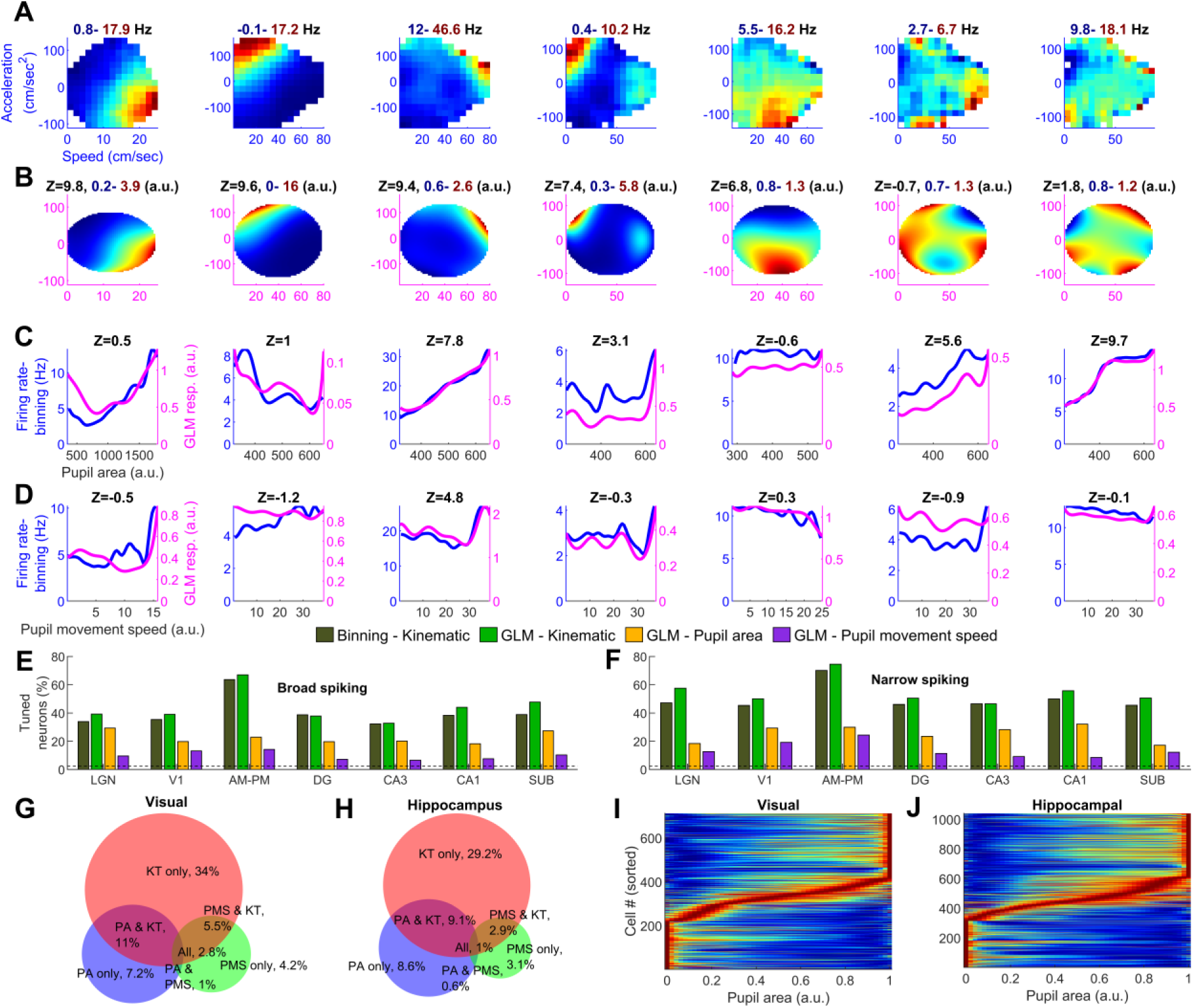
Kinematic tuning is more prevalent than pupil area tuning in all brain regions. **(A)** Seven representative cells with varing degrees of kinematic tuning computed using binning method. All but the last two cells had significant kinematic tuning, similar to Fig. 1. **(B)** Simultaneous estimate of the neurons’ (in A) modulation by the 2D speed-acceleration space **(C)** pupil size and **(D)** pupil movement speed (see *Methods* for data inclusion criteria). Similar to Fig. 1D, z-score of tuning and the range of modulation are indicated above the panels. For comparison, the binning method based estimates of pupil size and speed modulation are shown (blue) along with the GLM estimates (magenta). The two methods give comparable results, thus confirming the key results. **(E)** Fraction of significantly tuned broad spiking cells based on binning (dark green) and GLM (light green) methods are not significantly different in LGN, DG and CA3 (*p*>0.11), whereas GLM estimated greater kinematic tuning for other brain regions (*p*<0.04). Kinematic tuning (based on GLM) was significantly greater than the prevalence of pupil area or movement speed tuning for all brain regions (*p*<2.3×10^-3^). **(F)** Similar to (E), for narrow spiking cells. GLM and binning estimates for kinematic tuning were not significantly different, except for CA1 (*p*=0.01), and kinematic tuning was greater than pupil area or speed tuning in all brain regions (*p*<1.3×10^-5^). **(G)** Venn diagram of the percentage of cells from visual areas tuned for kinematic space (KT), pupil area (PA) and pupil movement speed (PMS) and their joint tuning. **(H)** Similar to (G), for hippocampal cells. Amongst cells tuned for only one of the three variables, more than three times as many cells were tuned for the kinematic space compared to pupil area or pupil speed. **(I)** Pupil area tuning curves based on GLM estimates, for all tuned cells from the visual areas. Each cell’s tuning curve was normalized between 0-1 (blue to red) and sorted based on the location of their peak response. 30.6% of cells responded maximally at the smallest pupil area values (constriction), 41.4% preferred the largest pupil area (dilation), and the remaining (27.9%) preferred intermediate pupil sizes. **(J)** Similar to (I), for hippocampus. Most neurons were tuned to either the most constricted (30.1%) or the most dilated (41.1%) pupil area with only 28.8% tuned for all the intermediate values of the pupil size.

We next tested the accuracy of GLM based estimates using a cross-validation procedure (see *Methods*). Briefly, we artificially generated Poisson spike trains with predetermined tuning for either the kinematic (Fig. S6A) or the pupillary variables (Fig. S6B), or both (Fig. S6C). There was little leakage of pupillary selectivity to kinematic tuning (Fig. S6D), validating the hypothesis that the observed kinematic tuning in the experimental data was not due to pupillary variables. This further demonstrates that kinematic tuning was not an artefact of pupillary dynamics. Do kinematic variables indeed have a stronger impact on neural activity than pupillary variables? Is this similar across all brain areas, including LGN that is just one synapse away from the retina? To test these surprising possibilities, we used a multipronged approach.

First, we compared the tuning responses from the GLM method for kinematic and pupillary tuning. This revealed that 23.4% of cells from visual areas were tuned for pupil dilation, of which 7.4% were tuned for pupil dilation, without significant kinematic tuning (Fig. 6E, G). Interestingly, this prevalence was similar even in the hippocampal regions, with a total of 21.7% tuned for pupil dilation, and 9.2% tuned for pupil dilation without kinematic tuning. In comparison, 33.7% of cells from visual areas and 30.1% of hippocampal neurons had kinematic tuning without significant modulation by either of the pupillary variables. Very few neurons were tuned for pupil movement speed (3-4%), which was close to chance levels (∼2.5%), and hence not probed further. Thus, the kinematic tuning was not only independent of pupillary dynamics, but it was stronger than pupillary dynamics modulation in all brain areas, including LGN that is a synapse away from the retina.

Interestingly, pupil-size tuned neurons preferred either very small or very large pupil size, with very few neurons preferring the intermediate pupil size (Fig 6I, J). This is similar to the kinematic-field accumulation at very low or very high speed and acceleration but not intermediate speeds and low acceleration (Fig 3C). The observation that very similar fractions of neurons preferred largest (41.2%), smallest (30.4%) and intermediate (28.4%) pupil size requires further investigations. Equally surprising is the finding that similar distribution of pupillary modulation was found in visual as well as the hippocampal areas without any task or changes in visuo-spatial stimuli.

### Refinement of GLM method reveals independent contributions of speed, acceleration and pupil dilation

To confirm these intriguing findings, we investigated the independent contribution of each of these four key behavioral variables: running speed, acceleration, pupil area and pupil speed. To this end, we refined the GLM method where each of these four behavioral metrics were treated as a separate one-dimensional variable (Fig. S7A-D, see *Methods*). Unlike the above joint-GLM, where the speed and acceleration contributions were measured jointly in the 2D speed-acceleration space, the new method treats each behavioral variable as an isolated one-dimensional variable, i.e. on a more similar footing. The results were broadly consistent with those reported above (Fig. 1, 2, 3, 6), further confirming that the kinematic tuning was indeed much higher than pupillary modulation in all brain regions (Fig. S7F). Many neurons were conjunctive (15.3%), with significant modulation for kinematic as well as pupillary variables (13.1% conjunction expected by chance). Pure speed, acceleration or pupil dilation tuned neurons were also present (Fig. S7A-D). Across all brain regions, speed tuning was the strongest, but significantly correlated with the acceleration-tuning, even after factoring out the contribution of pupillary variables (Fig. S7E). This shows that neurons were indeed jointly modulated by speed and acceleration resulting in kinematic fields in the 2D speed-acceleration space. Kinematic modulation was much stronger than the pupillary variables (pupil area or pupil movement speed), with the largest difference for AM-PM (61.3% for kinematic vs. 33.6% for pupillary) and least for dentate gyrus (38.8% vs. 27.5%). Pupillary variables significantly modulated the activity of at least 20% of broad spiking neurons in not only visual but all four hippocampal regions too, with the largest fraction in LGN (34.7%) and least in CA3 (22.2%). These results remained relatively invariant across all areas, from LGN, just one synapse away from the retina to subiculum (32.2%), the farthest (Fig. S7G).

A remarkably similar trend was found in even putative interneurons (Fig. S7H). The preponderance of kinematic tuning over pupillary tuning was largest in subiculum (54.5% for kinematic vs 23.7% pupillary) and least in V1 (52.7% vs 43.1%). The fraction of kinematically tuned neurons in each region showed the least difference between broad and narrow spiking neurons for subiculum (53.3% vs. 54.5% resp.), and largest for the dentate gyrus (38.8% vs. 53.1%). Thus, unlike many other types of neural coding, excitatory neurons and interneurons had similar kinematic tuning, which exceeded pupillary modulation in all brain areas.

### Same cell tuning for speed and acceleration are anti-correlated

To confirm these results and understand the nature of speed and acceleration tuning at the level of individual cells, we fitted the spike time series with a linear model that used speed and acceleration as predictors (see *Methods*) and confirmed that this provided a reasonable fit (Fig. S8). Comparing the magnitudes of speed and acceleration coefficients revealed that speed generated larger modulation than acceleration. This method estimated the contribution of either speed or acceleration, one variable at a time, hence it could be biased by the correlations between speed and acceleration. To address this, we noted that the occupancies in the kinematic space were invariably disk-shaped. Hence, we fitted the kinematic responses with Zernike polynomials up to the 3^rd^ order (Fig. S9, see *Methods*) and confirmed that it provided an accurate estimate using cross-validation procedure (Fig. S9A). The results again showed a two-fold larger modulation of spiking by running speed than acceleration (Fig. S9E).

Interestingly, comparing the speed and acceleration coefficients for the same cell revealed an inverse relation across brain regions. Typically, neurons preferred either the 2^nd^ quadrant, (corresponding to negative correlation with speed and positive correlation with acceleration) or the 4^th^ quadrant (positive correlation with speed and negative correlation with deceleration). This effect was robust and showed the same trend with different analytical techniques –the coefficients in the Zernike polynomial fits (Fig. S9B), linear time series fitting (Fig. S8B), and the coefficients of the GLM estimates (Fig. S10). Hence, while speed is a stronger modulator of neural firing than acceleration, neurons’ speed and acceleration modulations are not independent of each other, further demonstrating the inseparable nature of kinematic modulation and the existence of kinematic fields.

## Discussion

Here we examined the single unit activity of putatively excitatory (n=11329) and inhibitory (3533) neurons from seven different visuo-hippocampal regions in head fixed mice running spontaneously on a treadmill next to a monocular gray screen in a dark room. There were no rewards, tasks or even visual stimuli, which ensured that the neural selectivity to kinematic variables was not due to these factors. Despite the lack of these neural selectivity drivers, across all seven visual and hippocampal regions, at least 31% (CA3) and up to 61% (higher visual areas AM&PM) of broad spiking neurons’ activity was significantly and jointly modulated by running speed and acceleration (Fig. 2). As a result, these responses formed kinematic fields in the 2D, speed-acceleration phase space. The kinematic fields aggregated in two parts of the kinematic space –low-speed, high-acceleration (run initiation) and high-speed, low-acceleration (maximum speed) (Fig. 3).

The prevalence (40.0% of all hippocampal cells, n=9529) and size (about a third of the kinematic space) of the kinematic fields was comparable to that of place fields in several parts of the hippocampal formation^54,83–85^. Our measurements are from the intermediate and ventral hippocampus and the fraction of kinematic tuned cells was larger than the fraction of place cells in the ventral CA1^86^. The fraction of kinematic tuned cells was comparable to, or greater than the fraction of dorsal CA1 cells tuned for several other important features such as visual cues^63,72^ (33-39%), the location of conspecifics^87,88^ (13-18%), reward^89^, or time^90,91^ (∼30%). The kinematic tuned cells were also more prevalent than the fraction of grid^92^ (15%) and speed^14^ (13%) cells in the medial entorhinal cortex. Thus, the kinematic fields constitute a novel and major category of neural tuning in the visuo-hippocampal circuit.

Speed tuning has been extensively studied in the hippocampus of freely behaving rodents^10,12,13,48^, and recently in the visual cortex^2,3,5,7,26–28,30^. But often, running speed also induced changes in visual cues or the spatial location, or reward expectancy, thus the neural modulation cannot be attributed solely to the kinematics of running. In contrast, in our studies, there was virtually no change in visual cues or spatial location due to running, ensuring the purely kinematic origins of our findings, and consistent with speed tuning^7^ and modulation of visual stimulus selectivity by running^2,3,26^ in V1. Our results are also in agreement with widespread movement-related responses in the cortex^8^ as well as across the brain^9^.

Kinematic tuning in the LGN was comparable to that in the primary visual cortex, unlike prior work showing weak or non-existent modulation of thalamic visual responses by locomotion^2^, but consistent with other reports^27^. Further, the acceleration-dependence of single unit activity in LGN or the entire visuo-hippocampal circuit has received little attention and kinematic fields in the visual hippocampal circuit were unknown.

Many studies reported that hippocampal pyramidal neurons are sharply tuned for space but weakly tuned for speed, but the converse for interneurons^12^ with both cell types preferring high speed than low speed^10,12,13^. We found the opposite effect after accounting for the contribution of mean firing rates –broad spiking, putatively excitatory cells have better kinematic selectivity than interneurons, in dentate gyrus, CA3, and CA1. Further, higher visual areas had the highest kinematic selectivity, while the brain regions farther away from the higher visual areas, at both ends of the visual hierarchy (LGN and subiculum) had the least, and intermediate regions had intermediate levels of selectivity. Additional experiments can determine the mechanisms generating these patterns.

The speed tuning in hippocampal regions varies widely between studies from 10% to 25% in pyramidal neurons^14,17^, and 40% to 80% in interneurons^17,73^. In contrast, we found that both pyramidal and interneurons showed significant and comparable kinematic modulation in all brain areas, and the degree of kinematic tuning was relatively consistent across mice (mean±sem, 43.9±13.8% tuned across sessions) and across brain areas (Fig. 4). This suggests that the separation of function across pyramidal and interneurons in the hippocampus could be the result of motion induced changes in the multisensory visuo-spatial stimuli, which can vary across experiments, while the purely kinematic signal is universal across cell types and brain regions.

This hypothesis is further supported by our finding that the kinematic tuning was slightly but significantly diminished when the large gray screen was replaced by a movie (Fig. S2), which induces selective activation of large fraction of neurons in all parts of this circuit^72^. In this case, the movie images changed independent of the self-motion cues, causing greater dissociation with self-motion cues compared to the gray screen condition. Since most neurons in the visual areas were movie-tuned, our findings suggest that most neurons in the visuo-hippocampal circuit simultaneously encoded both internally generated kinematic cues and externally imposed visual cues. The interaction between the two could degrade kinematic tuning if the two are inconsistent. Further supporting this hypothesis, the kinematic tuning curves during grey screen presentation were often weakly correlated (mean correlation across brain regions between 0.23 to 0.48) with that during movie presentation in all areas.

Unlike the visual cue encoding which was more prevalent in the visual than hippocampal areas^72^, the kinematic tuning was comparable throughout the visual-hippocampal circuit. Although kinematic tuning was present in all layers of the visual cortex (Fig. 2), deeper layers typically showed stronger tuning than superficial layers, unlike visual information encoding, e.g. direction selectivity to drifting gratings is largest in Layer 4, compared to L2/3 and L6^93^. Amongst the hippocampal regions, kinematic tuning was more comparable across sub-regions, but deeper layers of CA1 had slightly greater kinematic tuning than the superficial layers (Fig. 2).

Several lines of analyses confirmed that the same neuron invariably encoded both speed and acceleration and spiked in a restricted region of the phase space of speed-acceleration, to generate kinematic fields. Although visual cortical neurons are insensitive to the visual stimulus acceleration^94,95^, they not only show self-acceleration dependence, with larger activation during acceleration than deceleration.

Surprisingly, 21.7% of neurons, both excitatory (19.4%) and inhibitory (28.4%), in all hippocampal subfields were significantly modulated by the pupil size (Fig. 5, 6 and S7). The degree of pupil size tuning of a neuron was significantly correlated with its modulation by kinematic variables (Fig. S7), despite any visual, vestibular or spatial contribution, indicating purely motoric mechanisms. Most tuned cells preferred either the largest or smallest pupil size, characterized by ramp-up or ramp-down responses; preferred tuning for intermediate pupil sizes was less prevalent (Fig. 5 and 6).

These pupil size modulation patterns were remarkably similar across all visuo-hippocampal areas –from LGN, just one synapse away from the retina to the subiculum, the farthest removed. The kinematic modulation far exceeded pupil size modulation in all areas, further demonstrating their ubiquitous nature. All analysis here were restricted to running epochs to avoid changes in gross behavioral state. The influence of pupil size (which is correlated with arousal^80,82,96^) might be greater during immobility since the kinematic behavior is absent.

Most dorsal CA1 neurons running in a virtual reality spiked as a function of the distance traveled by rats, and not the allocentric spatial location, in a variety of conditions and tasks, and independent of the specific visual cues^47,64,70^. Further, some CA1 cells fired as a function of the distance traveled by head-fixed mice during spontaneous bouts of motion in darkness without any visual stimuli or rewards^71^. Several analyses showed that the kinematic tuning could not arise due to such distance coding (Fig. S11, S12). For example, we found comparable kinematic tuning in all brain regions investigated, not just area CA1 where the distance tuning was reported, including the visual areas where distance tuning is unlikely to exist during virtual navigation due to constantly changing visual cues. On the other hand, it is plausible that kinematic tuning, when temporally integrated, could form the basis of distance coding to facilitate path-integration.

In sum, kinematic fields were widespread across the entire visuo-cortical system. Their prevalence and properties were not only similar across these seven brain regions, from the LGN to subiculum, but also across putative excitatory and inhibitory neurons. The same was true for pupil size tuning. Kinematic tuning exceeded pupil size modulation in all brain areas (Fig. 6). Similar to retinotopic space encoding in visual cortical areas and allocentric space encoding in hippocampal areas, the kinematic fields described here encode the self-motion space of running speed and acceleration. The prevalence and similarity of kinematic fields across the entire visuo-hippocampal circuit suggests that they are much more wide-spread and might be found in other sensory regions as well but arise from the motor cortices^16,97^. These kinematic fields could bridge the gap between ego-centric and allocentric spaces by providing self-motion signal in cortex as well as hippocampus for a variety of tasks including motoric interoception, self-motion tracking, and path integration.

## Acknowledgments

We thank the Allen Brain Institute for provision of the dataset, Dr. Josh Siegle and Dr. Jerome Lecoq for help with the dataset, Dr. Massimo Scanziani for input and feedback. C.S.P. is a Fellow of The Jane Coffin Childs Fund for Medical Research.

## Competing interests

Authors declare that they have no competing interests.

## Data availability

All data is publicly available at the Allen Brain Observatory – Neuropixels Visual Coding dataset (© 2019 Allen Institute, https://portal.brain-map.org/explore/circuits/visual-coding-neuropixels).

## Code availability

All analyses were performed using custom-written code in MATLAB version R2020a. Codes written for analysis and visualization are available from the corresponding authors at reasonable request upon publication.

## Author contributions

C.S.P. performed the analyses with input from M.R.M. C.S.P. and M.R.M. wrote the manuscript.

## Methods

### Experimental and surgical procedures, spike sorting and behavior

We used neural data and behavioral recordings made publicly available by the Allen Brain Institute at (https://portal.brain-map.org/explore/circuits/visual-coding-neuropixels). The details of the experimental pipeline and related publications are available at the above link^62^. Briefly, prior to implantation with Neuropixel probes, mice passively viewed a dataset including drifting gratings, Gabor patches and a natural movie and blank screen (spontaneous epochs) a couple of times. Neural signal obtained from probes was split into two channels. Data from the spiking band, sampled at 30 kHz with a 500Hz high pass filter was used. Head-fixed mice were free to run on a tilted running wheel. Their running speed, obtained from the running wheel movement, was synced with the neural data. Videos of eye were obtained at 30Hz and synced with the neural data and running behavior using a photodiode. For our analysis, the running behavior, measured at 30Hz was first smoothed with a Gaussian with σ=2 bins (66ms). Next, we computed the temporal derivative of the speed signal to obtain acceleration. The resultant signal was again smoothed with a Gaussian with σ=2 bins (66ms). The spontaneous running block consisted of 30 minutes where the screen had a uniform gray texture (blank screen). To eliminate the biasing effects of extensive immobility on kinematic tuning analysis, we restricted our analysis to running epochs from this 30-minute block. These were identified as epochs when a heavily smoothed (σ=660ms) running speed exceeded 2cm/sec. These running epochs were extended on either side for 5 seconds to capture very low speed and acceleration data. This ensured that relevant running behavior was fully captured while avoiding long epochs of immobility, which could signal a different brain state such as sleep. Pupil area was computed from the height and width of the ellipse fitted to eye tracking data. The center of the fitted ellipse was estimated at 30Hz, and the movement speed of this center, termed pupil movement speed, was obtained by computing frame to frame derivative of distance moved and smoothed by a Gaussian of σ=66ms. Kinematic variables of speed and acceleration along with these two pupillary quantities were interpolated to 100Hz for further analysis.

Prior to spike sorting, spikes were offset corrected and median adjusted to center the signal on zero. Spike sorting was automated using Kilosort2^9^, and the output was post-processed to detect unphysiological waveforms, and these were removed. The cleaned-up data was used for our analysis. *Neuropixel* probes were registered to a common co-ordinate framework^65^. Each recorded unit was assigned to a recording channel corresponding to its maximum spike amplitude and the electrode location assigned to a specific brain region using anatomical tracing. For statistical rigor, only units with a mean rate above 0.5Hz were used for analysis.

### Single unit categorization

Single units were classified into excitatory and inhibitory neurons using three different methods.

<u>Method 1</u>. Units with an average spike width (the duration between the minimum to subsequent maximum amplitude) greater than 0.45ms (but below 1.5ms, to prevent spurious, unphysiological spikes) were classified as broad spiking, putatively excitatory neurons. Putative inhibitory neurons were classified as narrow spiking, with spike widths below 0.4ms. This criterion was solely used throughout the paper, except for Fig S3, where additional criteria were tested to confirm results based on this method.

<u>Method 2.</u> Complex spiking index (CSI), which quantifies the reduction in spike amplitude of putative pyramidal cells during burst spiking^75^ was computed for all cells (Fig. S3), as done commonly for hippocampal place cells^47,74^. It was defined as

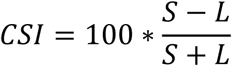

where S is the number of spike pairs with inter-spike interval < 20ms and amplitude of second spike less than the first. L is the number of spike pairs with inter-spike interval < 20ms and amplitude of second spike was greater than the first. Units with CSI greater than 20 were classified as putatively pyramidal (excitatory). Similarly, putative inhibitory interneurons cells were characterized to have CSI below 10. Units with intermediate CSI were not used, to make a conservative estimate.

<u>Method 3.</u> In addition to the CSI classification, we imposed the condition that excitatory cells should have a mean firing rate below 10Hz, and inhibitory interneurons should have a rate above 10Hz. Units which did not meet both the CSI and firing rate criteria were discarded, to obtain more conservative identification of excitatory and inhibitory cell types. Prevalence of kinematic tuning based on these different, more restrictive criteria is reported in Fig. S3.

### Kinematic tuning quantification

Procedures similar to those described previously were used^63^. The 100Hz sampled running speed and acceleration obtained as above, and spike counts from individual neurons, measured with 10ms bins were registered. Running speed and acceleration were binned using a 2-dimensional, linearly spaced speed-acceleration histogram, using 15 bins each. The data from extremes of acceleration, deceleration, and speed, denoted as the values beyond the extreme 0.5 percentile of data, were not used to avoid biasing by outliers. Additionally, bins with less than 500ms occupancy were discarded. This occupancy matrix was used to compute the number of spikes per unit time, i.e. firing rate, in each bin for each individual neuron under, resulting in the 2D kinematic tuning response. This was smoothed with a 2D Gaussian window of 3×3 bins. As commonly done for 2D hippocampal place cell firing rate maps^64^, the strength of modulation of kinematic fields was quantified as sparsity *s* of the rate-map where *r_n_* is the firing rate corresponding to the 𝑛𝑡ℎ bin, out of the *N=15×15=225* bins of the kinematic space:

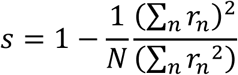

To assess the statistical significance of sparsity, we used a bootstrapping procedure^63,79^, which does not assume a normal distribution. Briefly, for each cell, spike train was randomly shifted by different amounts with respect to the kinematic behavior and the sparsity of the randomized data computed. This procedure was repeated 100 times with different values of random shifts. The mean value and standard deviation of the sparsity of randomized data was used to compute the z-scored sparsity of actual data using the function *zscore* in MATLAB. The observed sparsity was considered statistically significant if the z-scored sparsity of the observed spike train was greater 2, which corresponds to *p*<0.0228 in a one tailed t-test. A similar method was used to quantify significance of pupil size and movement speed in the GLM analysis (see below).

### Quantification of single cell and ensemble kinematic field properties

For each neuron that was significantly selective for kinematic space, we characterized its kinematic field using the peak firing location and field size in the speed-acceleration space, similar to the methods used to for quantifying 2D place fields^64,79^. To avoid artefactual biases by spurious high firing rate in a single bin, the peak of kinematic field-peak was defined as the firing rate weighted average of all bins with mean kinematic response above a threshold. This threshold was set as the 90^th^ percentile across all bins in the speed-acceleration space.

To quantify the size of the kinematic fields, a threshold based on the 90^th^ percentile from the corresponding shuffle data was used. Kinematic field size was defined as the percentage of bins having firing rate exceeding this shuffle-based threshold. These metrics are reported in Fig. 3 for broad spiking cells and repeated for narrow spiking units in Fig. 4.

Population responses for different brain regions were quantified using the kinematic rate-maps across all tuned units. To enable pooling and averaging responses across different sessions, the occupancy in kinematic space for each session was normalized between 0 to 1 for speed and-1 to 1 for acceleration. For this analysis, we focused on sessions which had at least 10% occupancy in each quadrant of the kinematic space to ensure similarity of occupancy across sessions. 14 out of the 18 sessions met these criteria. The same subset of sessions was also used for GLM estimation (Fig 6), and Zernike polynomial reconstruction (Fig. S9).

### Zernike polynomial reconstruction

We obtained Zernike coefficients for kinematic responses using the *zernike_recreation* function in MATLAB^98^, which provides the least square fit coefficients between Zernike polynomials and a given kinematic response. The first 6 polynomials, corresponding to orders 0, 1 & 2, were used. The coefficients corresponding to polynomials denoting variation along x-axis (speed) and y-axis (acceleration) were normalized by the coefficient corresponding to the 0^th^ order (constant) and compared in Fig S9.

### Linear dependence of spike trains on speed and acceleration

To assess the potential linear dependence of firing rate on speed and acceleration, we fitted the time series of each neuron’s firing rate with a linear model (Fig S8). To allow comparison between data with widely different range of speed and accelerations, we z-scored the speed and acceleration. The spike train of each neuron was fitted with these z-scored behavioral time-series. The *fitlm* function in MATLAB was used to perform linear fitting and obtain coefficients corresponding to constant *c*, speed coefficient α, and acceleration coefficient β. The coefficients α and β were divided by the constant *c*, to remove the effect of mean firing rates, and enable comparison across different neurons in Fig. S8.

### Comparison of excitatory and inhibitory kinematic tuning after spike down sampling procedure

The mean firing rates of narrow spiking cells were typically higher than that for broad spiking units. These high firing rates could artificially boost the kinematic tuning prevalence in narrow spiking units^82^. To address this, we focused on the broad and narrow spiking units with mean rates above 1Hz and randomly removed spikes such that the effective mean firing rate of 1Hz was obtained for each cell. Kinematic tuning as described above was repeated on these “down sampled” spike trains. This method ensured that the resultant kinematic tuning prevalence was compared between broad and narrow spiking units irrespective of their mean firing rates and is reported in Fig. 4 and S3.

### Kinematic tuning comparison between gray screen and movie presentation

To compare tuning during visual cue presentation, we used the experimental epochs when a 30 second clip from an isoluminant, black-and white, silent movie (Touch of Evil, 1958) was shown on the screen. Two blocks of 30 repetitions each were presented (total 30 minutes), one before and one after the gray screen presentation to enable robust comparison of kinematic fields. To quantify the similarity of kinematic tuning between the two visual stimuli conditions (movie vs. blank), we computed the correlation between the kinematic responses of the same cell in different conditions. Shuffle data were obtained by randomly reassigning the kinematic tuning curves across simultaneously recorded neurons between the two conditions. To ensure statistical rigor, only 11 out of the 18 sessions were used for this analysis, where the kinematic behavior was comparable during the two experimental epochs (correlation of kinematic occupancy map > 0.9) and all four quadrants of kinematic space had at least 10% of total time occupancy.

### Kinematic tuning prediction from pupil area and pupil movement speed

The pupil dynamics (pupil size and movement speed) were weakly (∼0.1 correlation, see Fig. S4), but significantly correlated with the kinematic variables. To rule out the possibility that kinematic tuning was solely caused by a leakage from the pupillary modulation of neural firing, we developed the following analysis. We first computed the average firing rate of neurons as a function of pupil area. Using this average firing rate, we predicted the instantaneous firing rate every 10ms, based on the observed pupil area at that moment. We then computed kinematic tuning, using this instantaneous, *predicted* firing rate that was solely based on the pupil area. This predicted kinematic response was compared with experimentally observed kinematic tuning in Fig. S5. A similar method was used to rule out pupil movement speed.

### Generalized Linear Model (GLM) based estimation of independent contributions of kinematic and pupillary variables on neural firing

We developed a Generalized Linear Model to estimate simultaneous and independent contribution of pupil dynamics and kinematic variables on spiking, using the *glmfit* function in MATLAB, as described previously^70,79^. Mice make fewer saccadic movements during running where our analysis is focused. These brief, ballistic movements are difficult to measure precisely with low frame-rate cameras (30Hz). Hence, we excluded potential saccades by removing top 0.5 percentile of pupil movement speed data which helps to prevent overfitting based on outliers. To ensure a good fit, extreme 0.5 percentile data from acceleration, pupil area and speed were ignored, and the top 0.5 percentile of running speed was also ignored, as above.

We developed two different GLM methods. In the first model, the running speed and acceleration were treated together as 2D kinematic space. The time-varying firing rate of each unit was modelled as an inhomogeneous Poisson process as a function of kinematic space and pupillary variables. Fourteen Zernike polynomials (orders 0 to 4) on the unit disk were used to model the kinematic space (speed-acceleration space). Pupil area and pupil movement speed were each modelled with sinusoids up to 5^th^ order, including a half-cycle sinusoid to allow non-periodic, continuously increasing or decreasing firing response estimation. We then used the estimated parameters from the best fit model to reconstruct the modulation of the firing rate by kinematic and pupillary covariates. The statistical significance of the resulting tuning curves was estimated by computing sparsity and a bootstrapping method described above using 100 shuffles for comparison with the binning estimates in Fig. 6. See (ref^79^) for more details and mathematical formulation.

Since kinematic space is two dimensional, while pupil area and pupil speed are each one-dimensional, the above method can potentially provide biased results. Hence, we developed another GLM model, where all 4 variables (running speed, running acceleration, pupil area and pupil movement speed) were each treated as independent one dimensional variables (All 1D GLM, Fig. S7). These were modelled with sinusoids upto the 5^th^ order as above. Tests of significance were performed as described above.

### Synthetic neuron analysis for GLM

To evaluate the extent to which correlations between kinematic and pupillary metrics can lead to spurious selectivity, we generated synthetic spike trains and performed GLM analysis using the first GLM model above. Synthetic spike trains were generated with mean rates ranging from 0.5 to 25Hz, and a range of firing between 0 to 50Hz. The synthetic neurons were generated with pre-determined input functions. These input functions were either a monotonically increasing, decreasing or uniform function of pupil area. Similarly for pupil movement speed. Synthetic kinematic fields were created with Gaussian functions centered at 9 different points in the speed-acceleration space. These average firing rates were then used to generate synthetic spike trains, using experimentally measured behavioral data, with the *poissrnd* function in MATLAB. These synthetic spike trains were fitted with the same GLM as described above to obtain estimates of kinematic and pupillary tuning curves. The resulting tuning curves were compared with the input functions to cross validate the GLM procedure (Fig S6 A-C). To directly estimate the potential leakage of pupillary tuning to kinematic tuning (and vice versa), we focused on the synthetic spike trains generated to have only pupil area or pupil movement tuning, but no kinematic tuning (or vice versa). The GLM results confirmed that the resultant spike trains did not have significant kinematic tuning (and vice versa, see Fig. S6).

### Binning based estimates of speed, acceleration, pupil area and pupil movement speed

To further confirm the GLM based results, we computed the tuning responses for all four behavioral variables using binning methods. The occupancy of behavior and spiking responses were binned, and outlier data were removed as described above. The resulting tuning curves were compared with the GLM based tuning curves in Fig. 6. It is commonly believed that neural firing rates increase with running speed. To quantify the linear dependence of neural firing on running speed, we computed the Pearson correlation coefficient between the one-dimensional, GLM based tuning curve and running speed. This was designated as “Speed coeff.” (and similarly for acceleration and pupil area coefficients). A similar method was used to determine if firing rates increased with acceleration, or pupil size (Fig S10).

## Extended Data

**Figure S1.**
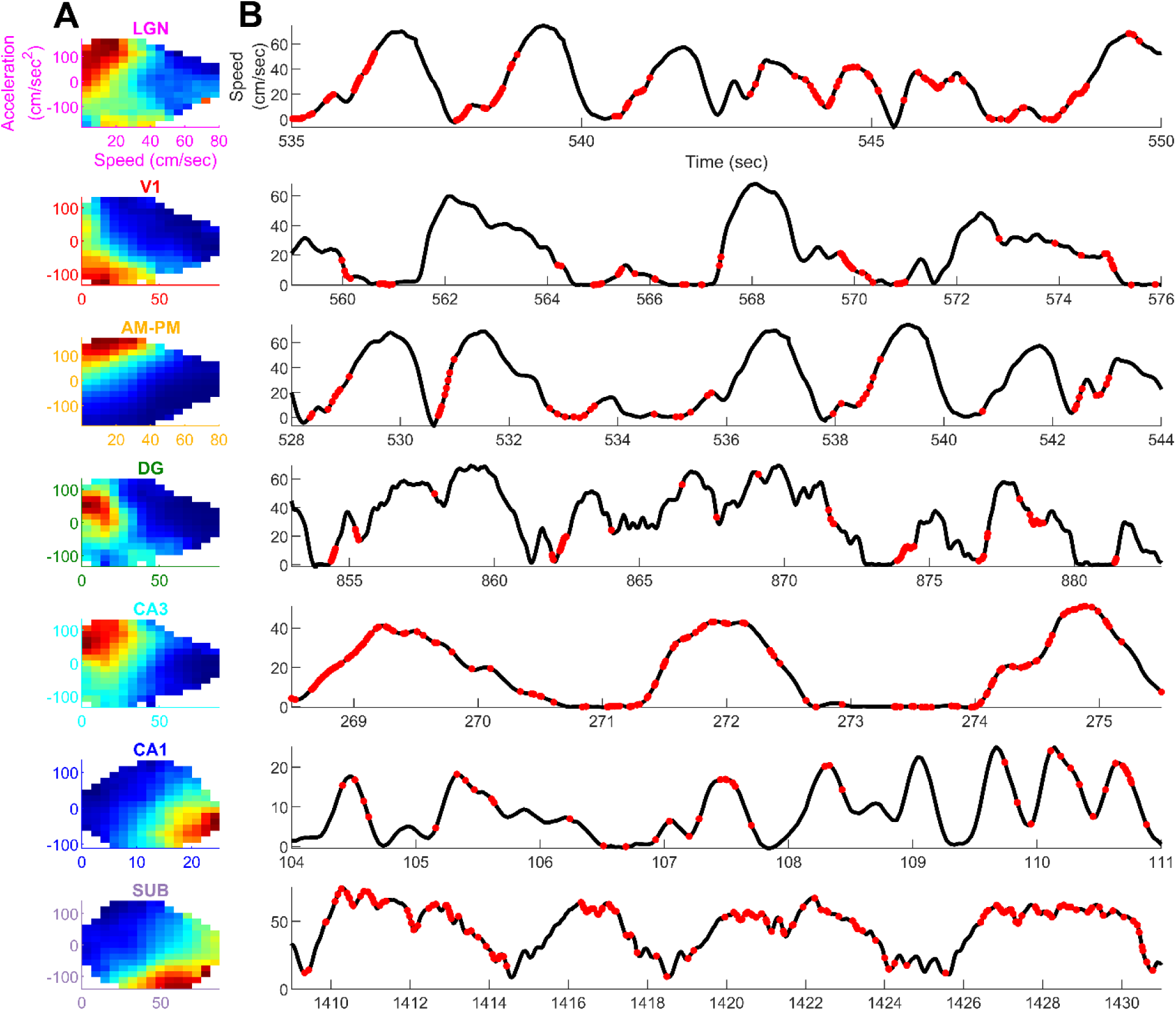
**Representative traces of kinematic tuning as a function of time (A)** One representative cell from each brain region, showing kinematic tuning (from Fig. 1). **(B)** A snippet of time, showing running speed in black, and spikes as red stars. Row 1,3 and 5 show neurons with preferential firing during epochs of increasing speed (acceleration), but relatively lesser spike during reducing speed epochs (i.e., deceleration), even at the same speed. Neuron #2 fires preferentially at low speed, but more so during deceleration, than acceleration. Neurons #6 & 7 fire preferentially at high speeds, but more so during decreasing speed epochs (deceleration) than acceleration.

**Figure S2.**
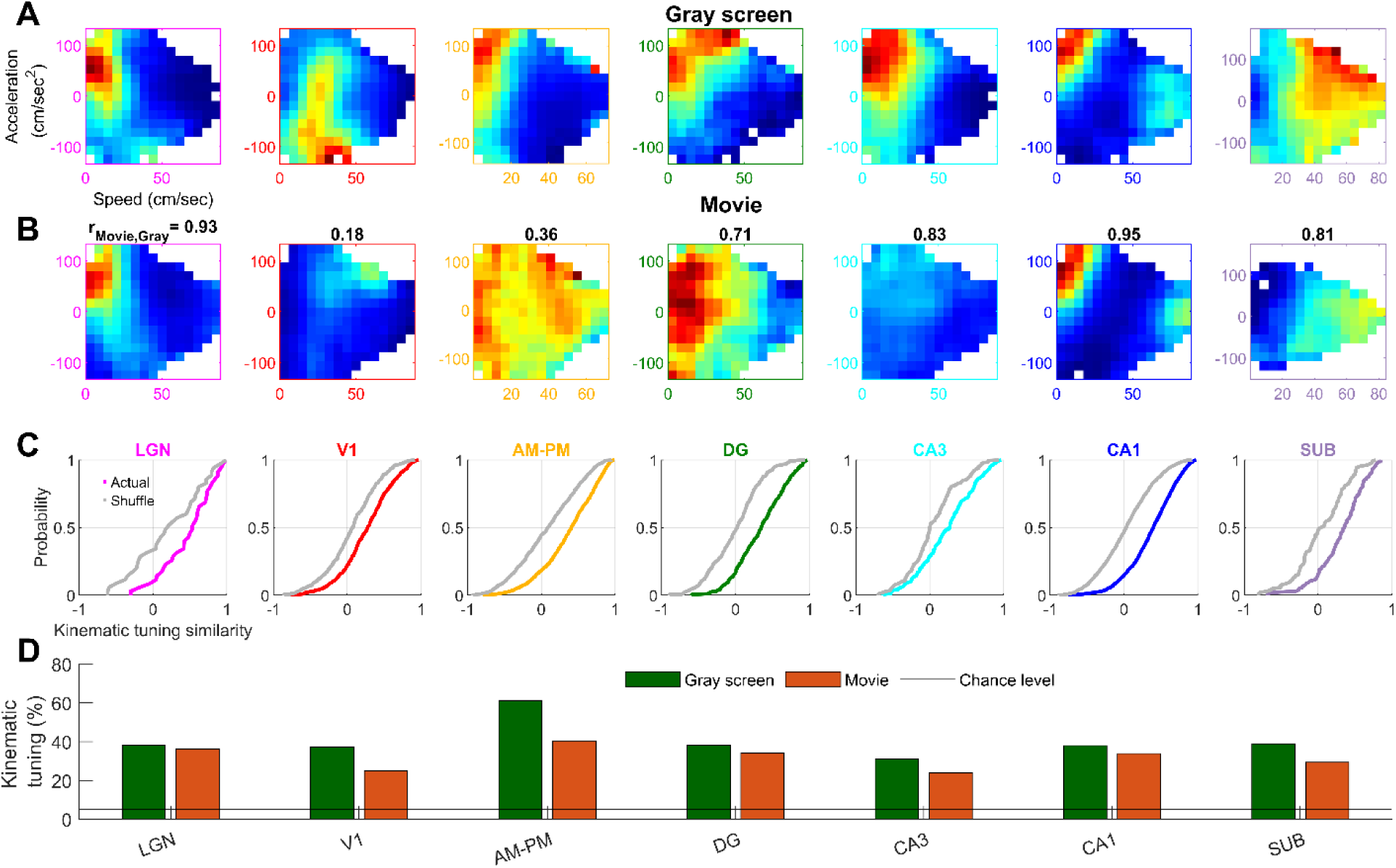
Kinematic tuning is intact but slightly weaker during movie presentation than that during blank screen. **(A)** A representative cell from each brain region showing kinematic field during gray screen presentation, and **(B)** during the presentation of a black-and-white, isoluminent, silent movie (see *Methods*). The correlation between the kinematic tuning during the two experimental epochs is indicated above. **(C)** Cumulative histogram of the similarity (correlation coefficient) between the kinematic tuning curves during gray screen and movie presentation for cells that were significantly tuned during either condition. Shuffle data (gray lines) were created by shuffling the neuron ID between movie and gray screen responses in the same session and same brain region. Kinematic tuning curves between the movie and blank screen conditions were significantly more correlated than that for shuffles in all brain regions (*p*<8.8×10^-5^). The mean values of actual correlations ranged from 0.23 in V1 to 0.48 in LGN, which is much larger than that for the shuffles (-0.0009 to 0.07). **(D)** Prevalence of kinematic tuning, (z-scored sparsity > 2) was above chance levels in all regions during movie presentation (red bars), with the minimum tuning in CA3 (24%) and maximum in CA1 (40%), and lesser than that during gray screen presentation (green bars) for all brain regions. The strength of tuning was significantly correlated across all brain regions (*r*>0.37, *p*<10^-31^ across the regions) between the two experimental conditions. Ratio of tuned cells (Movie/Blank) ranged between 0.66 for AM-PM, to 0.95 for LGN, and the strengths of tuning were significantly different for all brain regions (*p*<2.8×10^-3^), except LGN and DG.

**Figure S3.**
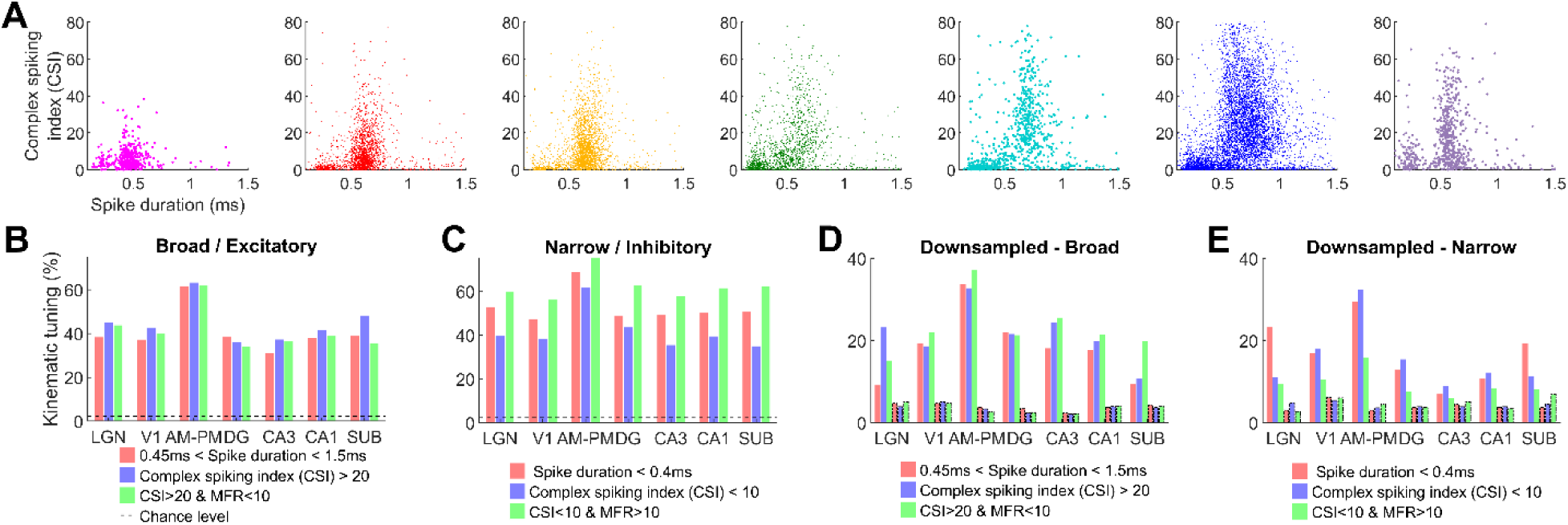
Different measures of separating putative excitatory and inhibitory yield similar results for kinematic tuning. **(A)** Spike duration (trough to peak distance of the average spike waveform) and complex spiking index shows two distinct clusters for all brain regions. As a result of this separation, broad (>0.45) and narrow (<0.4) spiking cells could be reliably separated. Units with wider spike waveforms typically had larger CSI. **(B)** Fraction of neurons with significant kinematic tuning, using spike duration or CSI metrics. All metric pairs resulted in similar tuning levels in all brain regions (*p*>0.05), except CA1 (*p*=0.02) and subiculum (*p*=0.005) when comparing spike duration vs. CSI based classification. Irrespective of the metric used, kinematic tuning obtained was above chance levels (black dotted line). **(C)** Similar to (B), fraction of tuned cells for putative interneurons or narrow spiking cells, classified using either spike waveform or CSI and firing rate-based metrics. Combined CSI and mean firing rate (CSI+MFR) criteria lead to significantly higher tuning prevalence than only CSI based classification (*p*<1.2×10^-4^) in all brain regions. Spike waveform duration-based classification also led to higher prevalence than CSI only classification (*p*<0.03). Similar to (B), kinematic tuning was above chance levels for all methods of classification tested. **(D)** To separate the effect of mean rates on kinematic selectivity, neurons with firing rates above 1Hz had their spike randomly down sampled to obtained kinematic tuning across neurons and brain regions at the fixed mean rate of 1Hz (see *Methods*). Even after down sampling, all brains regions for all metrics had significantly higher selectivity (*p*<0.01) in the broad spiking neuron population than chance (boxes with dotted outlines), except LGN with spike duration classification or CSI+MFR criteria (*p*>0.05). **(E)** Similar to (D), narrow spiking neurons’ kinematic tuning was reassessed by down sampling spike trains to 1Hz and showed significantly greater than chance selectivity prevalence in most brain regions and with most classification criteria. CA3 interneurons were not significantly different than chance level using spike duration or CSI+MFR criteria (*p*>0.3), and subiculum was not different than chance levels using CSI+MFR criteria (*p*=0.98). All other comparisons were significantly different between actual and chance tuning levels (*p*<0.04).

**Figure S4.**
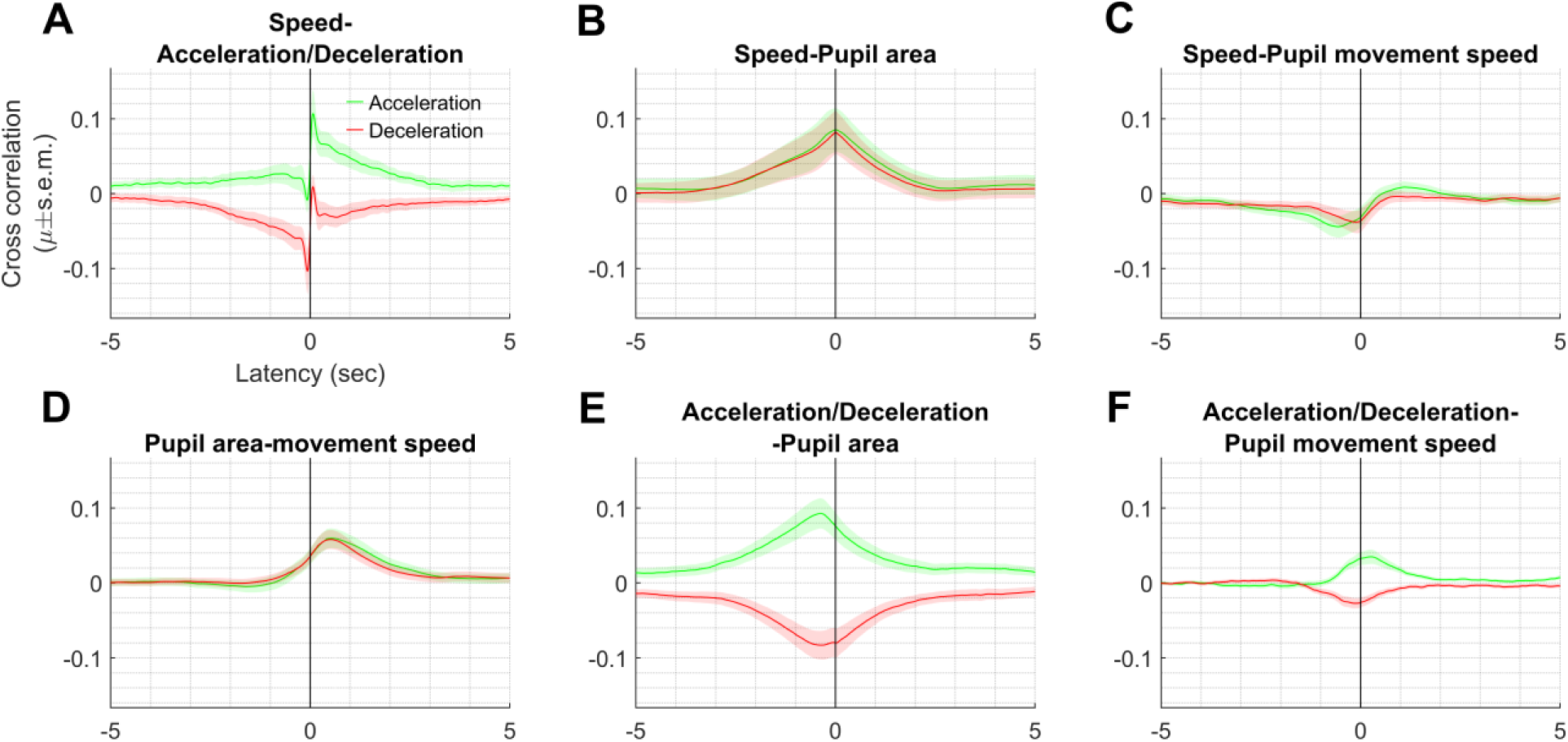
Kinematic behaviors were weakly correlated with pupillary behaviors. Across 18 sessions, **(A)** correlation between running speed and acceleration (deceleration) shows a peak (trough) such that acceleration precedes (follows) running speed by +70ms (-70ms). **(B)** Similar to (A), correlation between speed and pupil size (quantified by pupil area), had a maximum at zero latency suggesting speed and pupil area increase concurrently. **(C)** Running speed precedes pupil movement speed but it is maximally negatively correlated, indicating speed decreases before pupil movement speed starts increasing. **(D)** Pupil area increased after movement speed for accelerating as well as decelerating epochs, peak latency=530ms for accelerating, 520ms for decelerating epochs. **(E)** Acceleration peaked before pupil area increased, and also deceleration reduced (effectively more acceleration) before pupil area increases. Peak latency=-380ms and-350ms respectively. **(F)** Acceleration followed pupil movement speed (peak latency=260ms) and deceleration preceded it (peak latency=-130ms).

**Figure S5.**
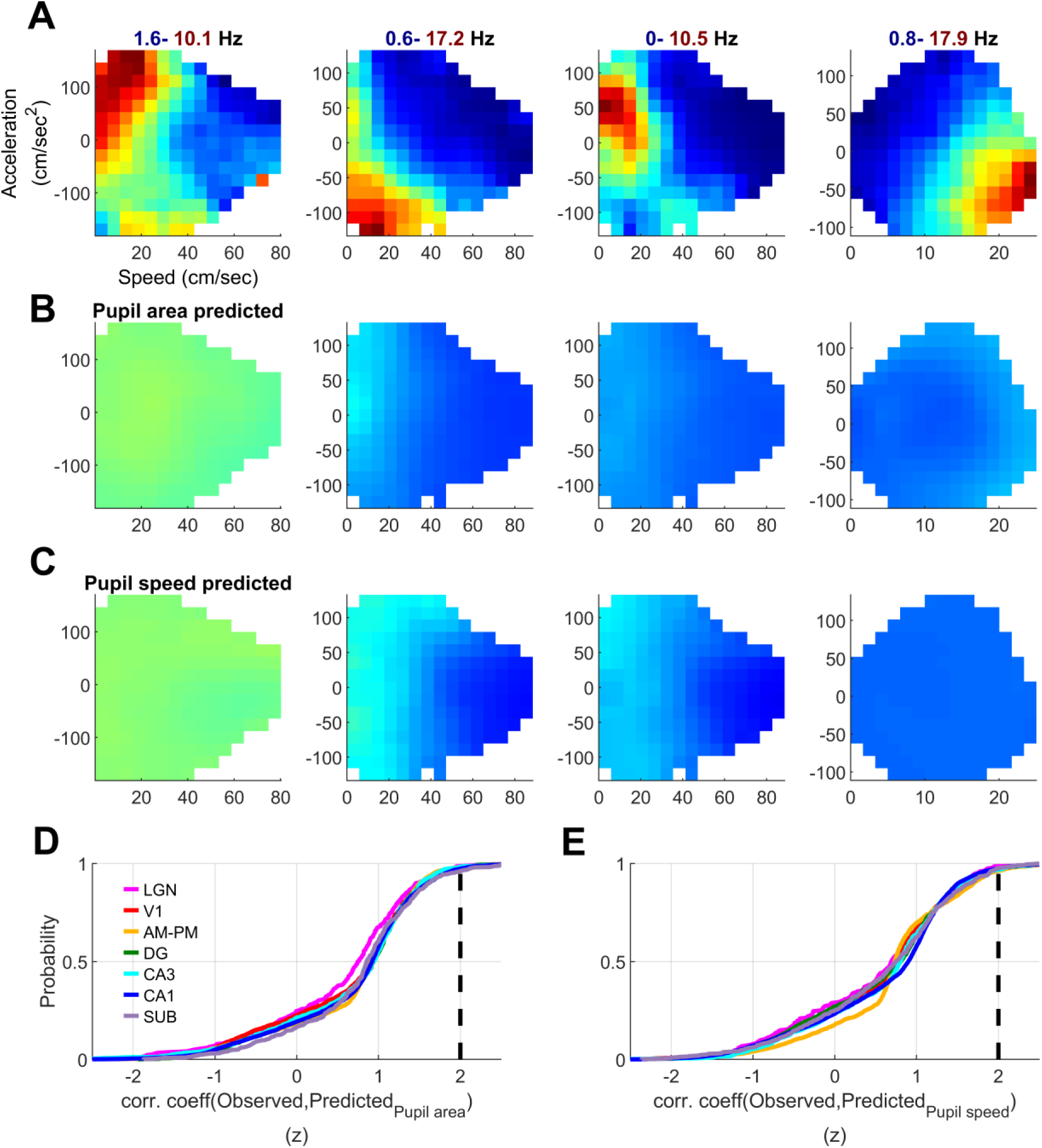
Kinematic tuning cannot be explained by pupil dilation or movement speed. **(A)** Four representative cells from Fig. 1 showing kinematic tuning. **(B)** Hypothetical kinematic tuning arising from correlated pupil dynamics for these cells was predicted using the average response of spiking to pupil dilation (quantified by pupil area), and the instantaneous relation between pupil area and running speed-acceleration (see *Methods*). Prediction of kinematic modulation using pupil area does not resemble the observed neural response in (A). **(C)** Similar to (B), kinematic tuning is not predicted by pupil movement speed. Panels through (A-C) for the same cell share color range as indicated above (A). **(D)** The relation between observed kinematic response and that predicted by pupil dilation was quantified by correlation coefficient and this was bootstrapped using mispairing between cells (see *Methods*). For all brain regions, less than 5% of correlations were significant at the z>2 level (black line), suggesting against an artefactual cause of kinematic tuning from pupil dilations/constrictions. **(E)** Similar to (D), less than 5% of correlations between actual kinematic tuning and that predicted by pupil speed were significant.

**Figure S6.**
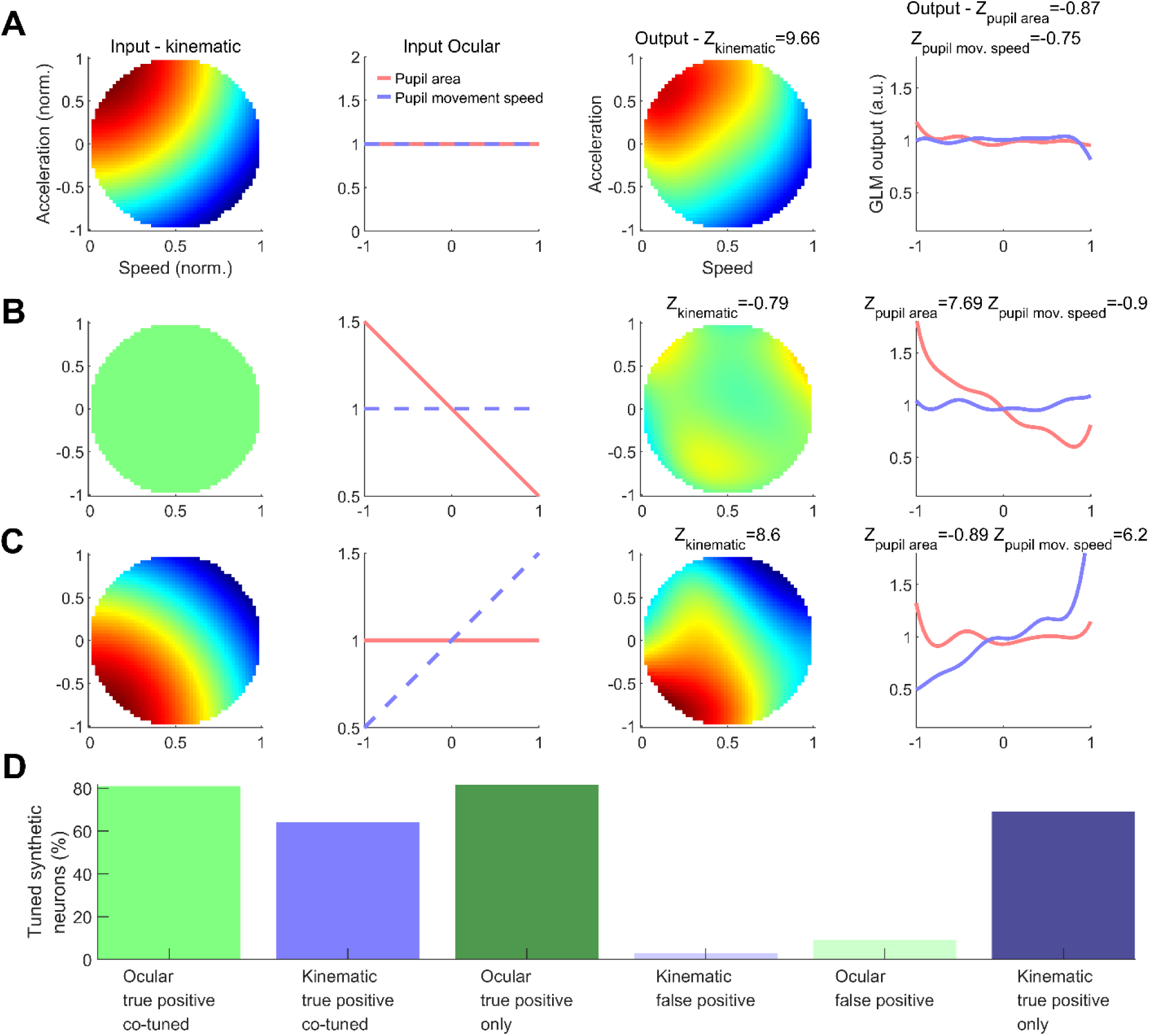
Simulation with Poisson spike trains validate GLM method and rules out artefactual leakage of pupillary tuning into kinematic tuning. **(A)** A simulated neuron that generated spikes according to a rate modulated Poisson process based on a predetermined kinematic tuning as input (preferred firing at top-left), with uniform probability of firing with respect to pupil dilation and movement speed. GLM faithfully recreated the tuning response of this neuron, without any leakage into the pupillary variables, thus cross validating the method. Resultant z-scored sparsity of tuning is indicated above. **(B)** Similar to (A), a neuron generating spikes only based on the pupil area selectivity and no kinematic tuning was rightly assigned high selectivity for pupil area. There was no leakage from the strong pupil area tuning into the kinematic tuning, which was effectively uniform. **(C)** Similar to (A), a neuron synthetically generated to spikes modulated by kinematic as well as pupil speed was rightly assigned high selectivity for pupil speed, as well as in the kinematic space. **(D)** Histogram of true and false positive rate from GLM across the simulated neurons. False positive rate of kinematic tuning, as a leakage from pupillary tuning was less than 5%. True positive rate for kinematic tuning was at 69.2% for neurons with only kinematic tuning and at 64.2% for synthetic neurons with kinematic as well as pupillary (pupil area or movement speed) tuning.

**Figure S7.**
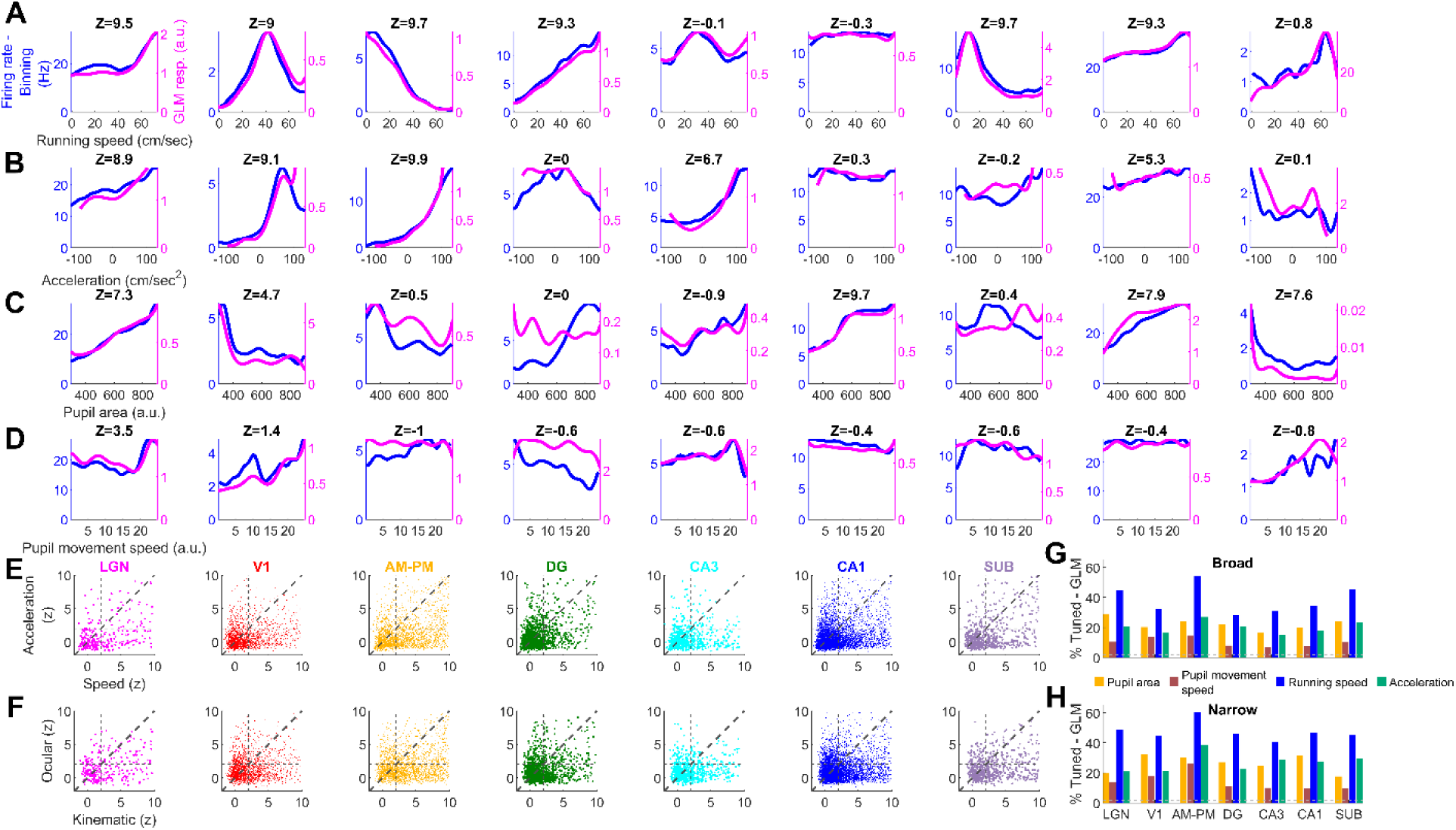
Kinematic variables were a stronger modulator of spiking than pupillary dynamics. **(A)** Nine representative cells showing similar neural response to running speed based on binning (blue) or GLM (magenta) analysis. This GLM analysis used kinematic as well as pupillary behavioral metrics as one-dimentional variables ensuring similar treatment and fair comparison between the 2 (see *Methods*). Z-score indicating strength of tuning is indicated on top. **(B)** Similar to (A), comparing acceleration responses. **(C)** Similar to (A), for pupil area. **(D)** Similar to (A) for pupil movement speed. The first cell showed tuned responses for all 4 variables, the second cell was tuned for speed, acceleration and pupil area but not pupil movement speed, the third cell was tuned for kinematic variables only and the next 4 cells were tuned for only 1 out of the 4 variables each. **(E)** Scatter plot for strength of tuning for speed and acceleration for all cells. Speed tuning was significantly greater (*p*<2×10^-7^) but also significantly correlated with acceleration tuning for all brain regions, even after factoring out the effect of mean firing rates (*r*>0.12, *p*<2.1×10^-3^). **(F)** Similar to (E), strength of tuning for kinematic variables (maximum of speed or acceleration tuning) was significantly greater (*p*<1.9×10^-5^) but also significantly correlated with tuning for pupillary variables (maximum of pupil area or movement speed), across all brain regions, even after factoring out the effect of mean firing rates (*r*>0.09, *p*<0.01). **(G)** Bar chart of percentage of tuned cells, showing highest tuning for running speed and least for pupil movement speed across all brain regions. Speed tuning was significantly greater than acceleration tuning in all brain regions (*p*<1.4×10^-7^), as well as pupil area tuning (*p*1.9×10^-5^). **(H)** Similar to (G), for narrow spiking cells, speed tuning was significantly greater than acceleration (*p*<0.03) as well as pupil area (*p*<1.3×10^-5^) tuning in all brain regions.

**Figure S8.**
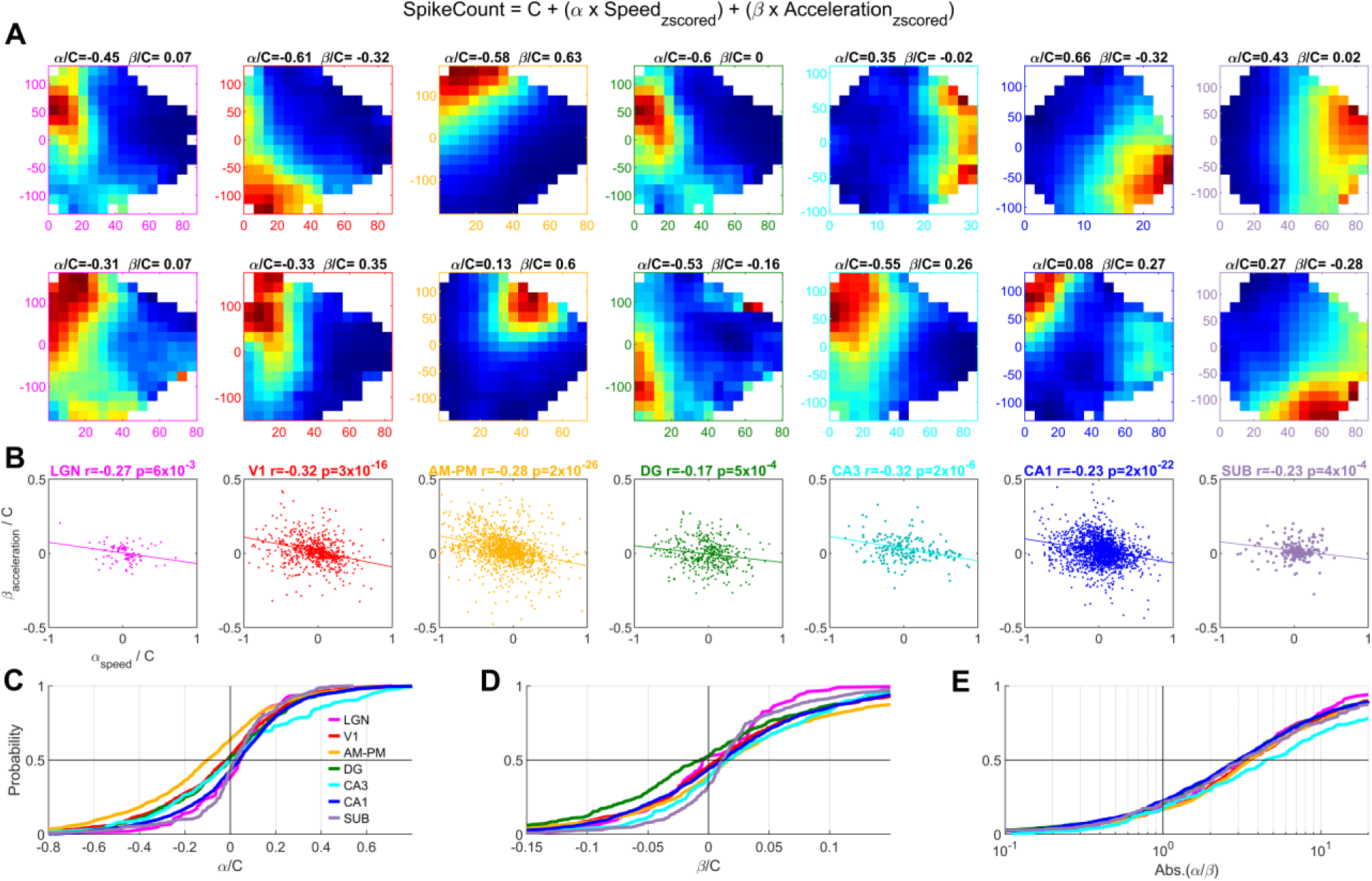
Negative correlation between speed and acceleration modulation of single units in all brain areas. **(A)** Fourteen representative cells showing kinematic tuning (from Fig. 1), with their coefficients of speed and acceleration from the linear model indicated above. For example, cell #1 (top, left) prefers low speed acceleration and has large negative value of α (normalized by the constant C from the model). This agrees with the kinematic response showing inverse relation between firing and running speed. The β (normalized by C) for this cell is ∼10 times lesser, in agreement with weaker modulation of firing with acceleration. **(B)** Speed and acceleration coefficients were significantly inversely related for all brain regions. Correlation coefficients and corresponding p-values are indicated above, for each brain region. **(C)** Cumulative histogram of the coefficient corresponding to running speed (normalized by the constant C). Largest bias of the coefficient was seen for higher visual areas, AM-PM (median=-0.1), indicating highest firing at low speeds. LGN, DG, CA3 and SUB distributions were not significantly different from having a mean of zero. **(D)** Similar to (C), for acceleration, showing small positive bias in all brain regions. This bias was significant in all brain regions (t-test for being centered at zero, *p*<2.3×10^-4^), except LGN and DG. **(E)** Cumulative histogram for absolute ratio of speed to acceleration coefficient, showing larger modulation with speed than acceleration based on the linear model (median ratio ranged from 3.0-DG to 4.5-CA1). This bias for a preference of speed over acceleration was significant for all brain regions (t-test for being centered at unity, *p*<5.3×10^-11^). This analysis was restricted to kinematically tuned, broad spiking cells.

**Figure S9.**
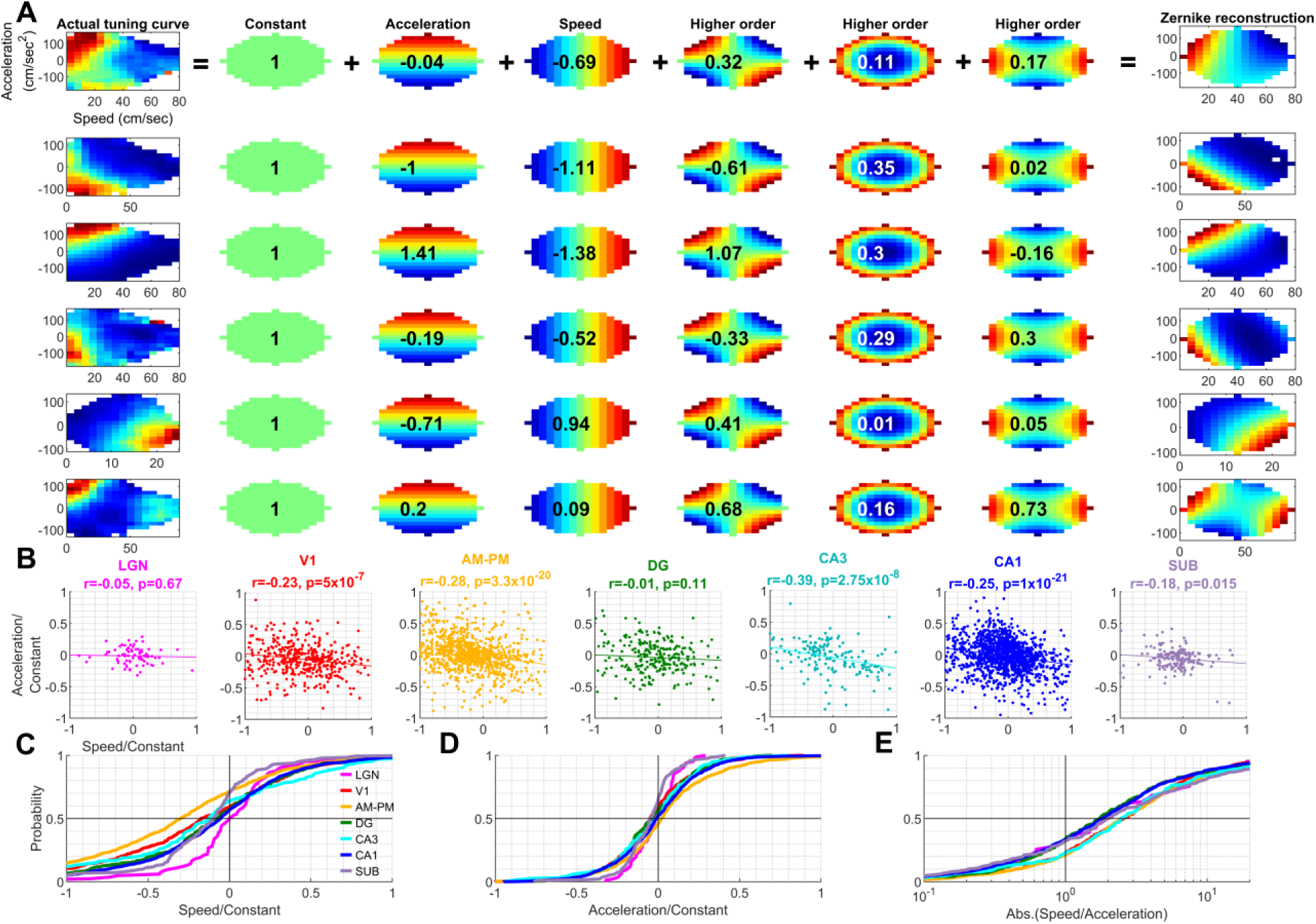
Zernike reconstruction of Kinematic tuning shows that most neurons were modulated by both speed and acceleration. **(A)** For six representative cells (from Fig. 1) the kinematic tuning was reconstructed using Zernike basis vectors up to the 3^rd^ order (see *Methods*). Coefficients corresponding to each Zernike basis vector are overlaid on top and normalized to the coefficient corresponding to the 0^th^ order (constant, 2^nd^ column). The coefficients corresponding to speed and acceleration were used for further quantification (see below). Note the similarity between actual kinematic tuning and that reconstructed by linear addition of these Zernike polynomials (last column). Speed coefficient is positive for the cell in the fifth row (since it has higher firing at higher speeds), whereas others are negative, as expected. **(B)** Scatter plot for the acceleration and speed coefficients (corresponding to the 2^nd^ and 3^rd^ Zernike polynomials, respectively), normalized by the constant term. Speed and acceleration coefficients were significantly negatively correlated for all brain regions except LGN and DG. The corresponding p-values indicated above. **(C)** Cumulative histogram of the speed coefficient (normalized by the constant term for each cell) for different brain regions. The speed coefficient showed the largest negative bias for AM-PM (median =-0.30) and was significantly (Wilcoxon signed rank test for zero mean, *p*<4.2×10^-3^) negatively biased for all regions except LGN. **(D)** Same as (C), but for the acceleration coefficient, which showed significant positive bias for V1 (*p*=1.2×10^-4^) and SUB (*p*=2×10^-6^), and significant negative bias for AM-PM (*p*=4.2×10^-4^). **(E)** The coefficient corresponding to speed was, on average, larger than that for acceleration by 1.7*x* (DG) to 2.5*x* (V1) fold. This analysis was restricted to kinematically tuned, broad spiking cells, from sessions with comparable kinematic space occupancy in all 4 quadrants (n=14 sessions, see *Methods*).

**Figure S10.**
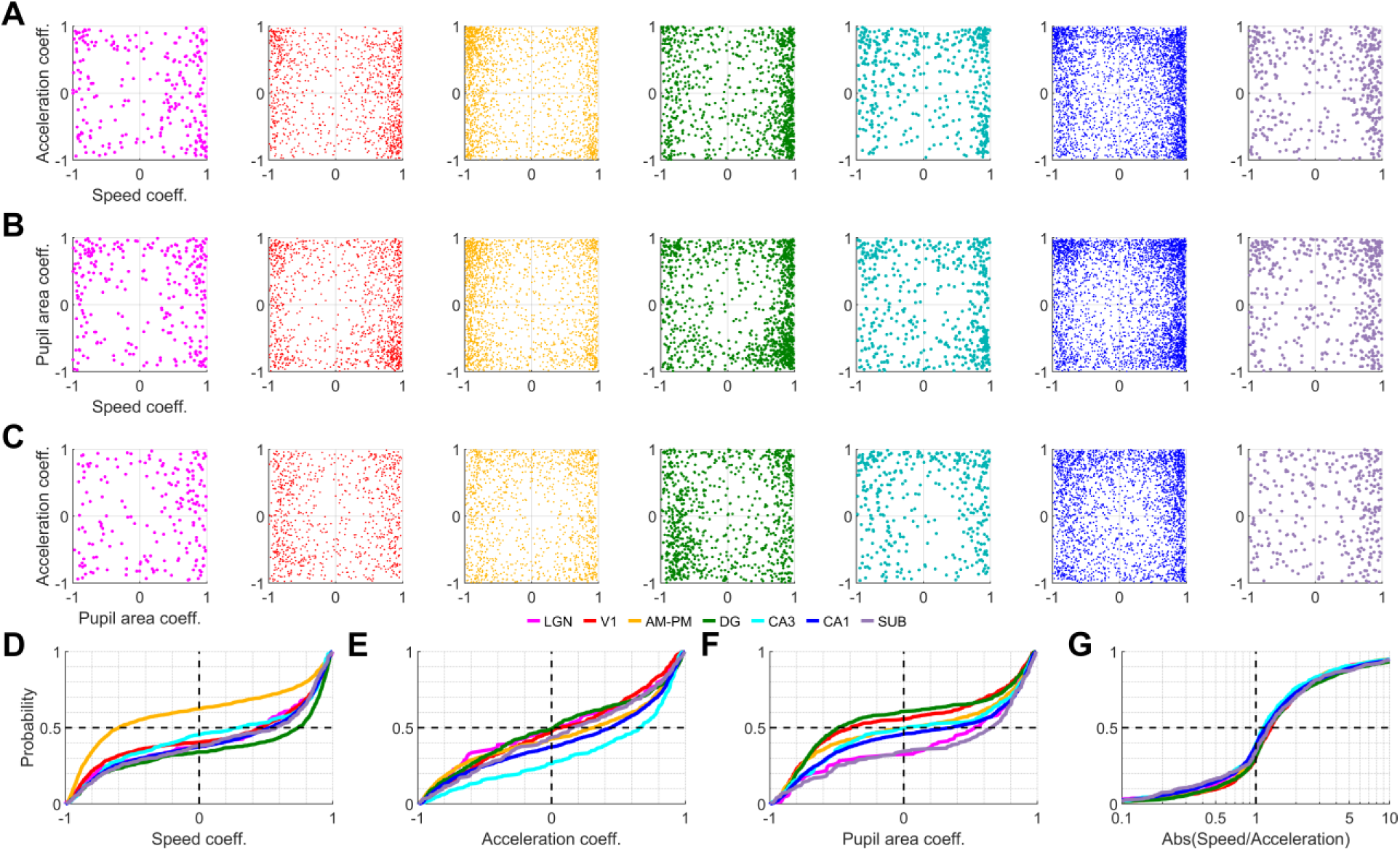
Spike rate-speed and rate-acceleration correlation coefficients are inversely correlated with each other. **(A)** Speed and acceleration coefficients (coeff.) were computed using the one-dimensional tuning estimates from GLM (see *Methods*). The speed and acceleration coefficients were inversely correlated and this relation was statistically significant for all brain regions (*r<*-0.13, *p*<0.02), except for subiculum. Cells with either speed or acceleration tuning were used for this analysis. **(B)** Same as (A), but for pupil area and speed coefficients. Speed and pupil area coefficients were significantly negatively correlated for all brain regions (*r<*-0.11, *p*<4.6×10^-4^), except LGN and SUB. **(C)** Same as (A), but for acceleration and pupil area coefficients, which were not significantly correlated for most brain regions, except DG (*r=*-0.14, *p*=7.3×10^-5^). **(D)** Cumulative histogram of speed coefficients; all brain regions had a bias towards positive values (median values for brain regions - +0.3 for CA3 to +0.73 for DG), except AM-PM (median =-0.6). All distributions were significantly different from having a mean of zero (t-test, *p*<8×10^-4^). **(E)** Same as (D), but for acceleration coefficients. Many distributions were significantly different than being centered at zero (p<0.01), except LGN, V1 and DG. The median values were positive and ranged between 0.01 for DG to 0.65 for CA3. **(F)** Same as (D), for pupil area coefficients. All distributions were significantly different than being centered at zero (t-test, p<0.01) except AM-PM and CA3 (t-test, p>0.05), but the median values ranged between (-0.49 for DG to +0.65 for SUB). All histograms were created with only those cells which were significantly tuned for the respective variable. **(G)** The cumulative distribution of the ratio between the speed and acceleration coefficients (using same subset of cells as (A)) were biased to values above 1 for all brain regions (t-test, *p*<2×10^-4^), with median values ranging between 1.14 for CA3 to 1.3 for V1.

**Figure S11.**
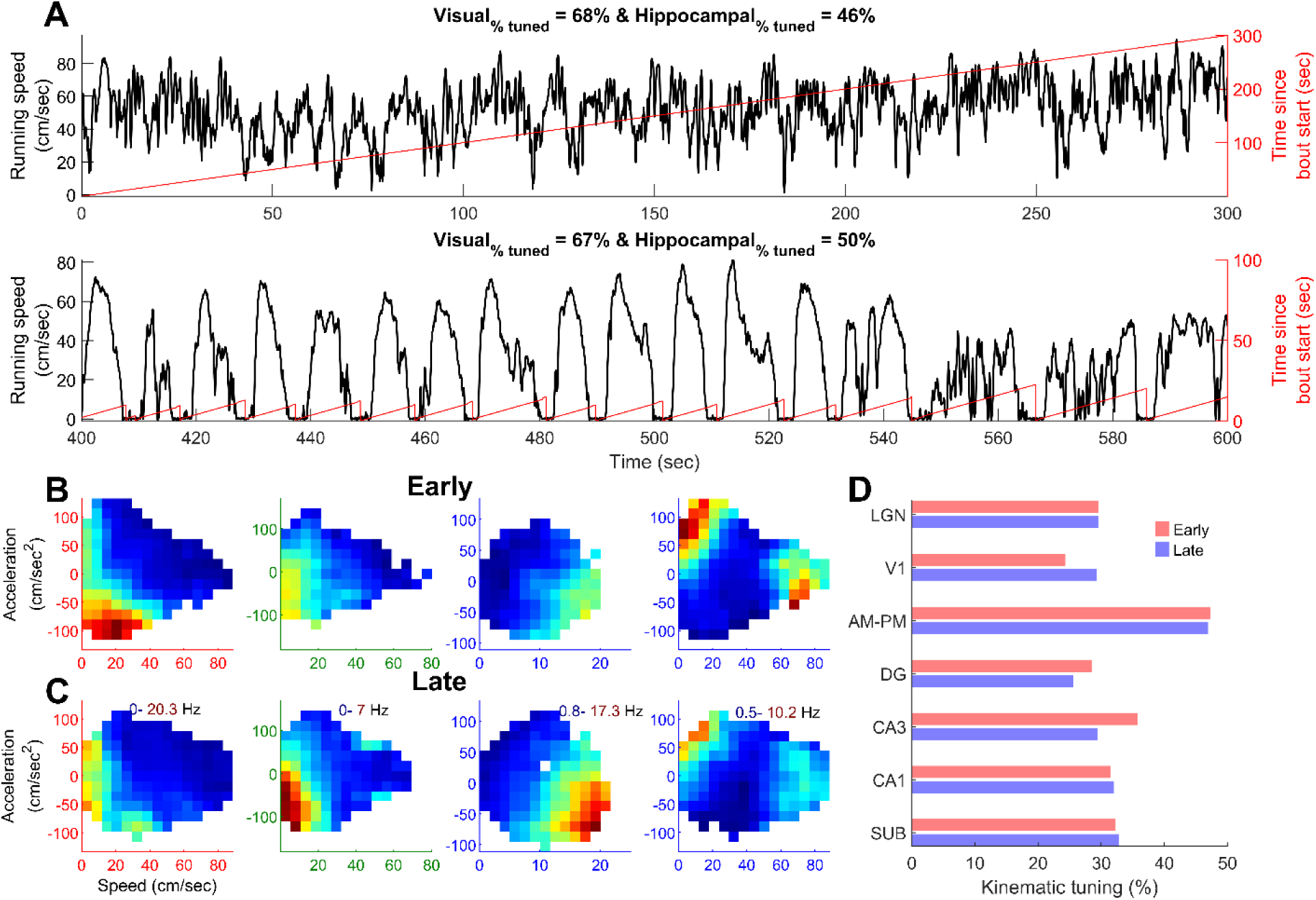
Kinematic tuning persisted during long as well as short bouts of running, inconsistent with a purely internally driven mechanism. To assess whether kinematic tuning was independent of the duration of continuous running, we split the epochs of running into “bouts”. A bout of running was defined as a segment of time starting from when the animal’s running speed exceeded 2cm/sec and continued till the speed dropped and stayed below 2cm/sec for more than 2 seconds. Across the 18 sessions, a wide variety of running bouts occurred. **(A)** An example of a long bout, where running speed (black, top) never dropped to zero. The time in the long bout is overlaid (red). In another session with multiple running bouts (bottom), comparable fraction of tuned cells were found in visual and hippocampal regions (indicated above). **(B)** To quantify this across all cells, we computed the kinematic selectivity of the same neuron to data from the early (within 20 seconds from start) and late parts of a running bout, in sessions with at least 3 minutes of data in each category (n=8 sessions). Kinematic tuning persisted in the early parts of the running bouts, as shown here for four example cells (V1-red, DG-green and two from CA1-blue). **(C)** For the same neurons as (B), the tuning curves during late parts of the running bouts were qualitatively similar as early part of the bout. Same color axes were used for the same cell across (B) and (C), and this range is indicated above. **(D)** Kinematic tuning was not significantly different in terms of the prevalence across broad spiking neurons (*p*>0.05) for all brain regions except V1 (*p*=0.03), where later parts of a running bout led to higher kinematic tuning. The same amount of data in each session for late and early parts of a running bout was used to rule out its contribution to selectivity. This resulted in less data for both subsets compared to the entire experiment resulting in the lower prevalence of kinematic tuning here compared to the full experiment (Fig 2).

**Figure S12.**
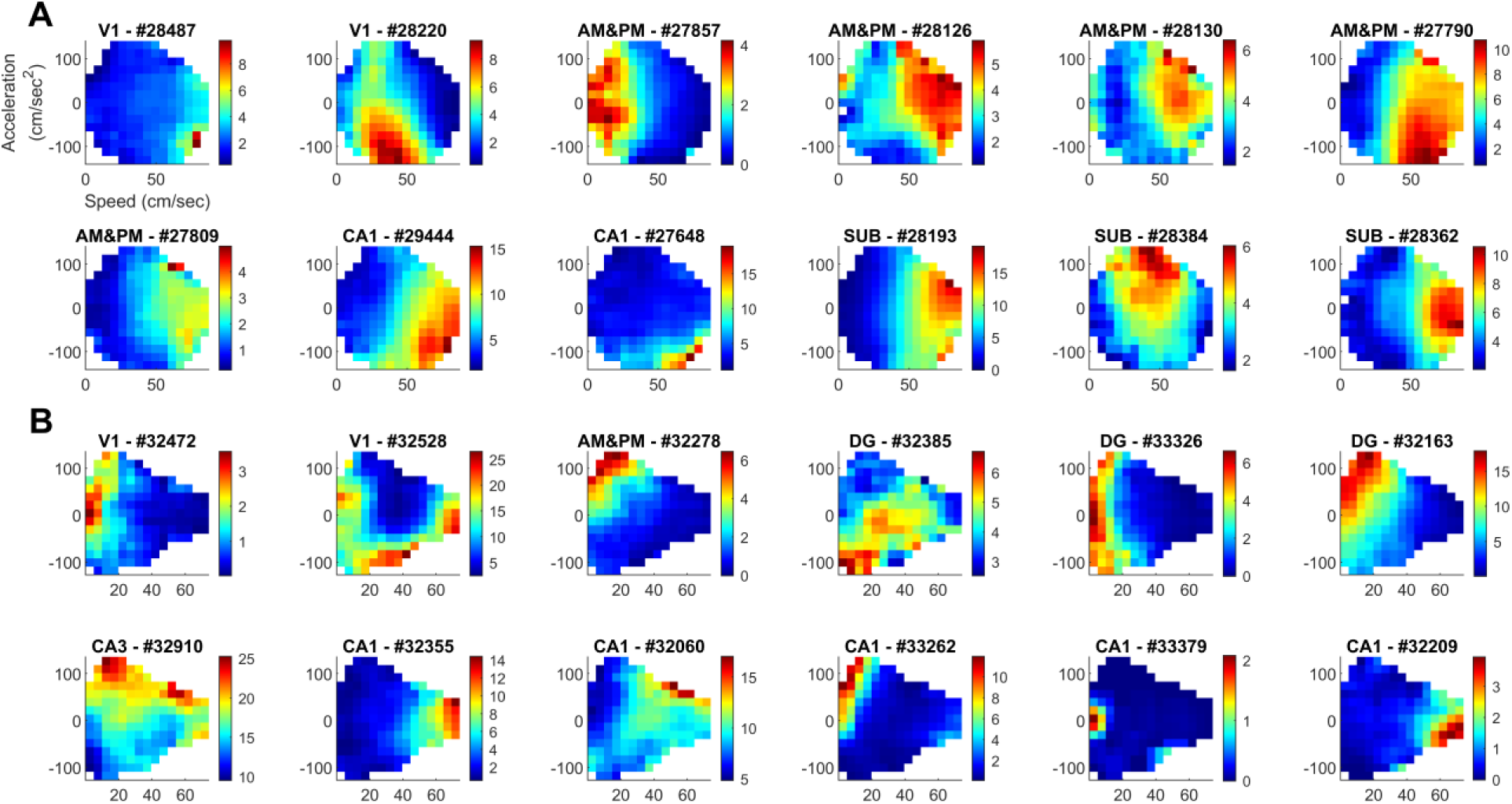
Simultaneously recorded cells within the same session show diverse kinematic tuning curves. **(A)** Twelve example cells from visual cortical as well as hippocampal regions recorded simultaneously in a session where the animal ran continuously (First example session from Fig. S11A). The firing rate range is indicated to the right. **(B)** Similar to (A), for a session where the animal stopped and resumed running behavior multiple times (2^nd^ example session from Fig. S11A).

## Notes

### Competing Interest Statement

The authors have declared no competing interest.

